# Development of a Bipyrimidineamide based α-Helix Mimetic Lead Compound for efficient Targeting of MDM2 in Triple-Negative Breast Cancer

**DOI:** 10.1101/2024.03.02.582899

**Authors:** Jasmin Linh On, Vitalij Woloschin, Franziska Gier, Jia-Wey Tu, Sanil Bhatia, Thomas Lenz, Andrea Kulik, Kai Stühler, Dieter Niederacher, Hans Neubauer, Tanja Fehm, Thomas Kurz, Knud Esser

## Abstract

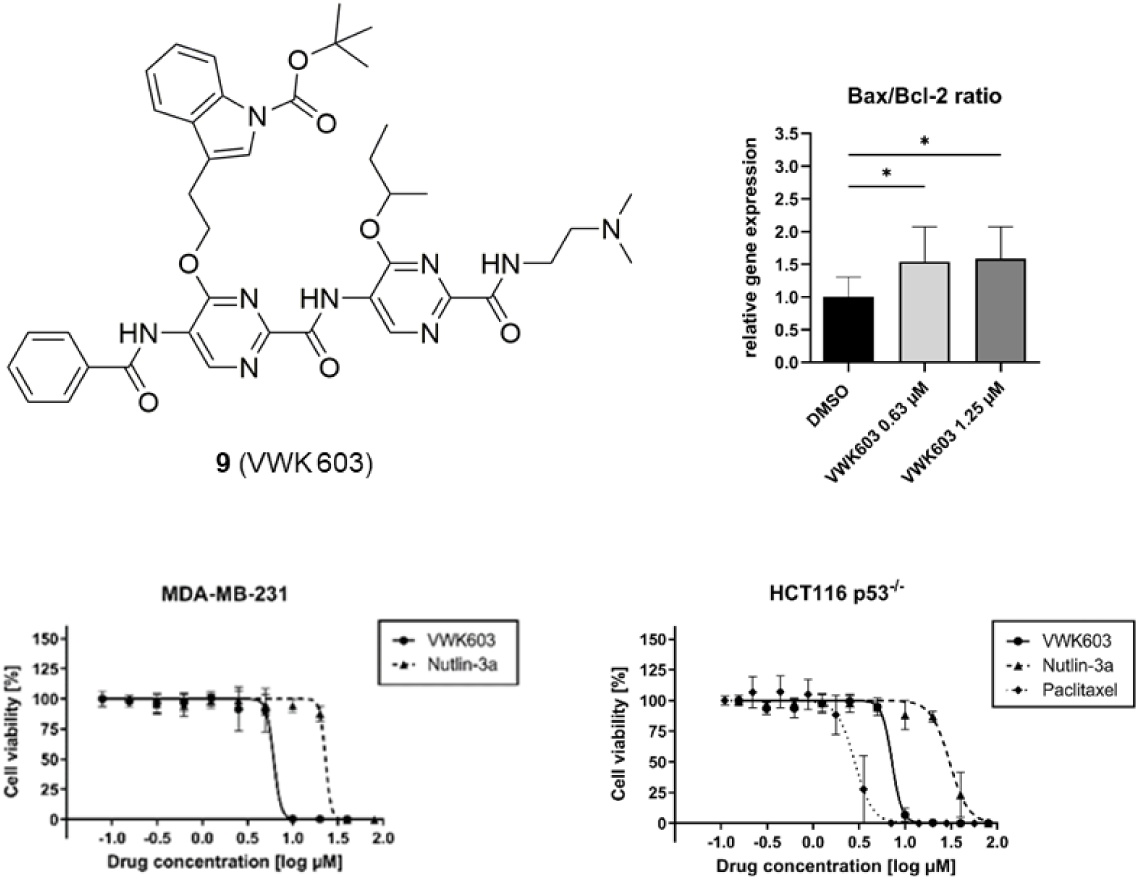

Triple-negative breast cancer (TNBC) represents the most aggressive form among breast carcinoma subtypes. Due to limited therapy options, identification of novel active pharmacological compounds is an urgent medical need. A promising approach in cancer treatment is the pharmacological inhibition of murine double minutes 2 (MDM2)-p53/p73 interactions inducing apoptosis in tumors. We here describe a novel bipyrimidineamide based α-helix mimetic **9** (VWK603) which was designed as a lead candidate to target MDM2. **9** (VWK603) potently induced cell death in the TNBC cell lines MDA-MB-231, MDA-MB-436 and MDA-MB-468 with IC_50_ values ranging between 3.7 µM and 6.6 µM. The anti-tumor activity was about four more potent higher than determined for the MDM2-specific inhibitor Nutlin-3a. Mechanistic analysis revealed induction of cellular apoptosis as the underlying mode of action of **9** (VWK603) anti-tumor activity. Since toxicity was observed to be reduced in non-cancerous breast cells, these studies make **9** (VWK603) a promising candidate for further preclinical MDM2 inhibitor development.

---

Triple-negative breast cancer (TNBC) accounts for 15 – 20% of all breast cancer subtypes and represents the most aggressive subtype. Due to the lack of cellular targets well known for other breast cancer subtypes, namely estrogen-, progesterone- and human epidermal growth factor 2 receptor, therapeutic options are limited. Additionally, the current TNBC treatment regimens are broadly based on (neo-)adjuvant chemotherapies associated with significant side effects and often leading to high chemoresistance.^1, 2^ Although with immunotherapeutic agents and PARP inhibitors first more targeted therapeutic options are available^1^, these drugs are often limited to distinct TNBC subgroups and regularly lead to therapy resistance development early after therapy initiation^2, 3^. Therefore, novel therapeutic strategies are urgently needed. However, the identification of new effective drugs in TNBC remains challenging.

To identify novel treatment options and to tag new therapeutic vulnerabilities in TNBC, we performed a drug screening assay using an in-house library of potential α-helix mimetics. We hereby identified the novel α-helix mimetic **9** (VWK603) which potently induced cell death in MDA-MB-231 cells. The bipyrimidineamide **9** (VWK603) had been developed to specifically target murine double minutes 2 (MDM2) and computational structural alignment studies stressed that **9** (VWK603) belongs to the MDM2 inhibitor class. MDM2 is an E3 ubiquitin ligase and the main negative regulator of the tumor suppressor protein p53 by inhibiting its transcriptional activity and promoting its proteasomal degradation.^4^ Pharmacological inhibition of MDM2-p53 protein-protein interaction leads to the activation of p53 and induction of cellular apoptosis or cell cycle arrest.^5, 6^ In addition to p53, MDM2, within the same region, has also been shown to interact with p73^7, 8^ resulting in pharmacological reactivation of p73 to induce apoptosis in tumor cells especially when p53 is not functional.^9–11^

Importantly, since a large number of key protein-protein interactions are mediated by α-helices^12, 13^, specifically interfering with α-helix interactions have been proven to be powerful therapeutic approaches in certain diseases including cancer.^14–16^ Since also the MDM2 protein-protein interactions have been shown to involve distinct exposed side-chain residues of particular α-helices^17^, the development of novel MDM2-specific α-helix mimetics has a high potential to improve current strategies for MDM2 inhibition. VWK603 was developed with the intention to mimic the hot spot amino acids Leu26, Trp23 and Phe19 of p53 binding in the deep hydrophobic cleft of MDM2. The *sec*-butyl residue mimics the side chain of Leu26, the *N*-Boc-(indol-3-yl)ethyl moiety mimics interactions of Trp23 and the benzoyl group mimics Phe19 of p53.

The synthesis of the bipyrimidineamide **9** (VWK603) was performed out from pyrimidine monomers according to Scheme 1. The reaction of azlactone **1** with the amidine hydrochloride **2** in the presence of triethylamine provided pyrimidone **3** as starting material in 51% yield.^18, 19^ A subsequent Mitsunobu reaction of **3** with the corresponding commercially available alcohols (butan-2-ol or tert-butyl 3-(2-hydroxyethyl)-1H-indole-1-carboxylate) in the presence of triphenyl phosphine and diisopropyl azodicarboxylate in tetrahydrofuran furnished the 4-alkoxypyrimidine **4** (45%) and 4-aralkoxypyrimidine **5** (65%) yield. The pyrimidine derivative **4** was converted into pyrimidincarboxamide **6** in an aminolysis reaction of its methyl ester moiety with *N,N*-dimethylethylenediamine in ethanol at room temperature in 79% yield. Next, the benzoyl protecting group of **6** was removed with a methanolic sodium hydroxide solution in dichloromethane at room temperature to provide aminopyrimidine **7** in 81% yield. The lithium carboxylate **8** was synthesized by alkaline hydrolysis of the methyl ester group of pyrimidine **5** using lithium hydroxide in a mixture of tetrahydrofuran and water in 86% yield. In a HATU mediated coupling reaction the lithium carboxylate **8** and the 5-aminopyrimidine **7** were converted to the bipyrimidineamide **9** in 29% yield. Upon purification, **9** was obtained with a chemical purity of 97.4%. Identity and purity of **9** were confirmed by ^1^H- and ^13^C-NMR-spectroscopy (Figure S11, S12) as well as by mass spectrometry (Figure S13).

**Scheme 1.**
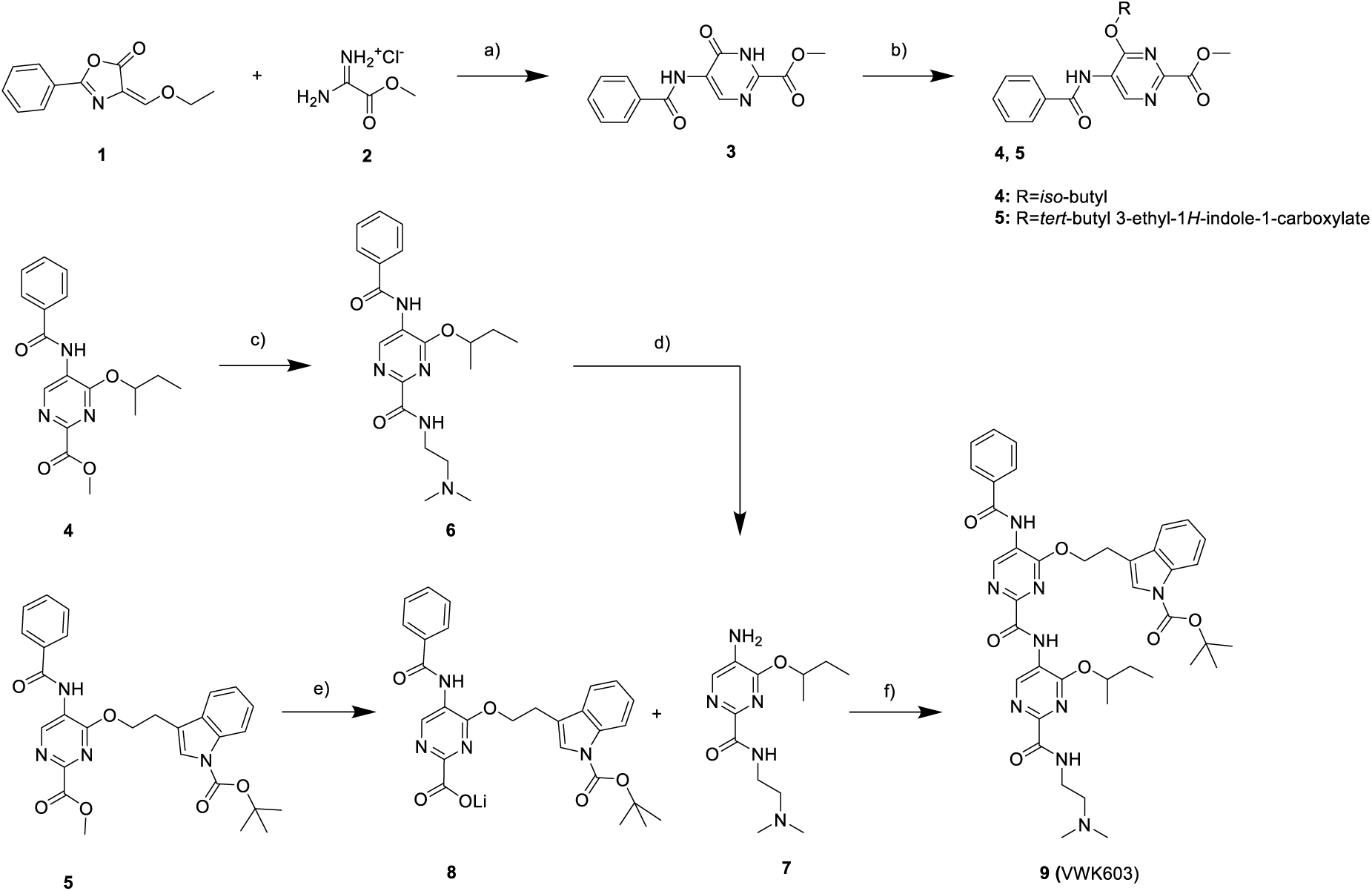
Synthesis of the bipyrimidinamide 9 (VWK603) from pyrimidin monomers. a) NEt_3_, acetonitrile, reflux, 3 h; b) PPh_3_, alcohol, DIAD, THF, 0 °C – r.t., 4 h; c) N,N-Dimethylethylenediamine, EtOH, r. t., 16 h; d) NaOH, dichloromethane:MeOH, r. t., 24 h. e) LiOH, THF:H_2_O, r. t., 24 h; f) HATU, DMF, r. t., 24 h.

The anti-tumor activity of **9** (VWK603) in MDA-MB-231 as initially discovered in our drug screening illustrating significant induction of cell death at 10 µM was first investigated in the TNBC cell lines MDA-MB-231, MDA-MB-436 and MDA-MB-468 in more detail. Cells were treated with different concentrations of the novel compound **9** (VWK603) or the reference compound Nutlin-3a, a well-characterized inhibitor of MDM2-p53 interaction.^20^ After performing cell viability assays, inhibitory dose-response curves were generated and IC_50_ values were calculated. Both compounds, **9** (VWK603) and Nutlin-3a, significantly reduced the cell viability in a dose-dependent manner in all TNBC cell lines (Figure 1). The IC_50_ values of **9** (VWK603) in MDA-MB-231, MDA-MB-436 and MDA-MB-468 were 6.58 µM, 5.93 µM and 3.67 µM, respectively, while the corresponding IC_50_ of Nutlin-3a with 23.15 µM, 20.84 µM and 19.36 µM resided in a marked higher range. These data illustrate that **9** (VWK603) efficiently targets cancer cell viability of TNBC cells and possesses an about 3.5-fold superior pharmacological potency than the reference compound Nutlin-3a.

**Figure 1.**
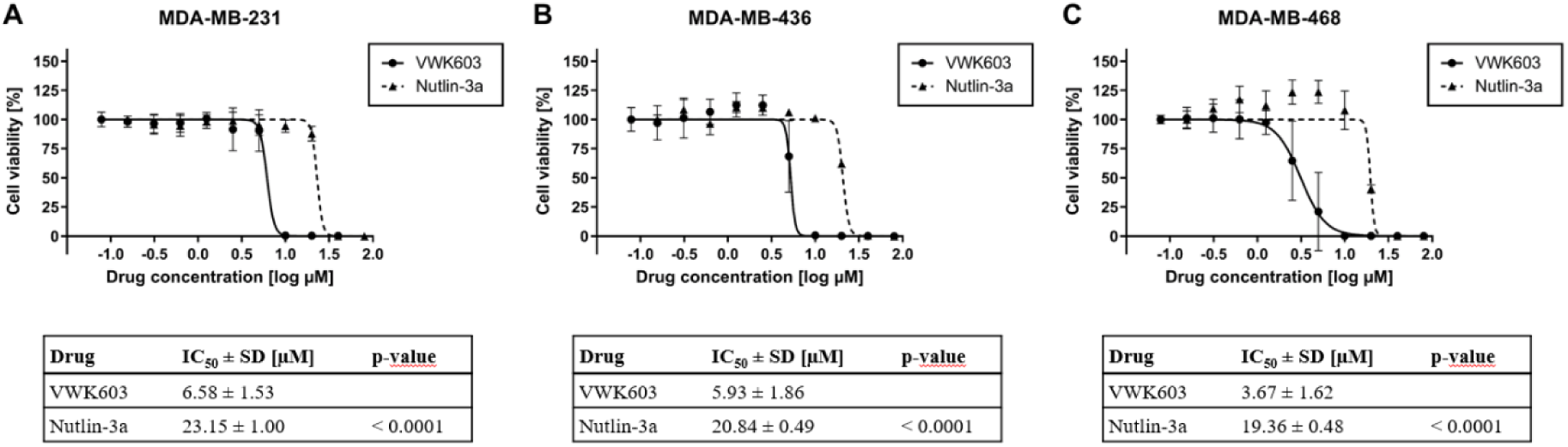
Analysis of cell viability in TNBC cell lines after VWK603 and Nutlin-3a incubation. MDA-MB-231 (A), MDA-MB-436 (B) and MDA-MB-468 (C) were seeded and incubated for 24 hours before treatment. **9** (VWK603) or Nutlin-3a was added at displayed concentrations and incubated for 72 hours. Treatment was repeated for another 72 hours. Subsequently, cell viability was determined by CellTiter Glo^®^ assay. The graph shows the mean ± SD n≥3. Table listed below the graph contains the summarized IC_50_ values ± SD. Statistical analysis was performed by using unpaired student’s t-test. P-values indicate the significant difference between the two compounds.

Importantly, the cell lines used in our studies, as most TNBCs, harbor a p53-inactivating mutation.^21^ In fact, MDM2 inhibitors have also shown to act via targeting MDM2-p73 interaction, especially when physiological p53-function is missing.^9, 10, 22^ Actually, a significant anti-tumor activity by MDM2 inhibitors has also been shown in TNBC cells by interfering with MDM2-p73 interaction.^11, 23, 24^ Since our investigation illustrated a very pronounced antitumoral effect of **9** (VWK603) in TNBC cells harboring an inactivating p53 mutation, we were interested to know whether the reduction of cancer cell viability occurs independently from cellular p53 expression like illustrated for other MDM2 inhibitors.^9, 22, 23, 25, 26^ To address this important point, we investigated the effect of **9** (VWK603) on cell viability in the colon carcinoma cells HCT116 p53^+/+^ and HCT116 p53^-/-^. Cells were treated with different concentrations of **9** (VWK603) or the reference MDM2 inhibitor Nutlin-3a. Additionally, Paclitaxel was used as control since this drug has been described to induce apoptosis independently from p53.^27^ After performing cell viability assays, inhibitory dose-response curves were generated and IC_50_ values were calculated. Both compounds, **9** (VWK603) and Nutlin-3a, and the chemotherapeutic agent Paclitaxel reduced the cell viability in a dose-dependent manner in both cell lines regardless of the p53 status (Figure 2). The corresponding IC_50_ values of **9** (VWK603) in HCT116 p53^+/+^ and in HCT116 p53^-/-^ were 6.64 µM and 7.52 µM, for Nutlin-3a 28.03 µM and 30.59 µM, and for Paclitaxel 1.25 µM and 2.82 µM. As reported in earlier studies, Paclitaxel significantly induced cell death in both cell lines.^27, 28^ For **9** (VWK603) and Nutlin-3a, no significant difference was observed regarding to the p53 status. Importantly, there was a marked difference in the IC_50_ values determined for **9** (VWK603) and Nutlin-3a of approximately 4-fold confirming the superior anti-tumor activity of **9** (VWK603) over Nutlin-3a discovered in TNBC cells also in colon carcinoma cells (Figure 2, Table 1).

**Figure 2.**
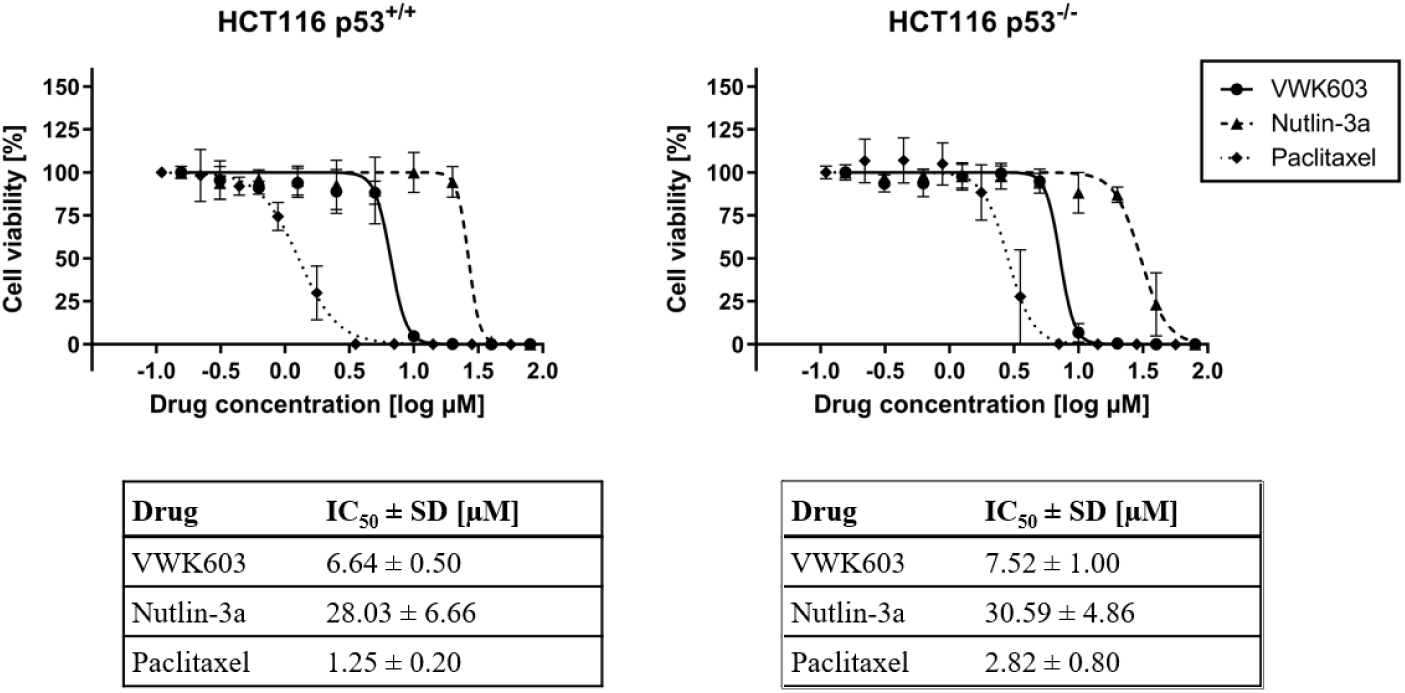
VWK603 effect on cell viability in HCT116 p53^+/+^ and HCT116 p53^-/-^ cells. HCT116 p53^+/+^ (left panel) and HCT116 p53^-/-^ cells (right panel) were seeded and incubated for 24 hours before treatment. **9** (VWK603), Nutlin-3a or Paclitaxel were added at displayed concentrations and incubated for 72 hours. Treatment was repeated for another 72 hours. Subsequently, cell viability was determined by CellTiter Glo^®^ assay. The graph shows the mean ± SD n≥3. Table listed below the graph contains the summarized IC_50_ values ± SD.

**Table 1.** Comparison of the pharmacological potency of compounds tested in HCT116 p53. ^+/+^ **and HCT116 p53**^-/-^ **cells**

| Compounds | P-value (HCT116 p53<sup>+/+</sup>) | P-value (HCT116 p53<sup>-/-</sup>) |
| --- | --- | --- |
| VWK603 vs. Nutlin-3a | < 0.0001 | < 0.0001 |
| VWK603 vs. Paclitaxel | < 0.05 | NS |
| Nutlin-3a vs. Paclitaxel | < 0.0001 | < 0.0001 |

P-values in the table summarizes the significance of the IC<sub>50</sub> difference between the compounds tested in HCT116 p53<sup>+/+</sup> (left) or HCT116 p53<sup>-/-</sup> cells (right) as illustrated in Figure 2. Statistical analysis was performed by using one-way ANOVA.

MDM2 inhibitors have widely been shown to induce cellular programmed cell death via apoptosis, in p53 wildtype as well as more recently in p53-mutated tumor cells.^9, 11, 23, 25, 26^ We therefore aimed to determine the underlying mechanism of cell death induction by **9** (VWK603) by quantifying the gene expression of the pro-apoptotic protein Bax and anti-apoptotic Bcl-2, which inversely regulate the cellular apoptosis via the mitochondrial way, and calculating the Bax/Bcl-2 ratio to quantify the apoptosis inducing effect. The Bax/Bcl-2 ratio determines the susceptibility of cells to apoptosis stimuli^29^ and is used to assess the apoptotic potential of novel identified compounds when applied to the cells in a non-toxic concentration.^30^ A Bax/Bcl-2 ratio < 1 is characteristic for resistant cells and a higher Bax/Bcl-2 ratio is characteristic for sensitive cells to an added apoptotic stimulus.^31^

MDA-MB-231 were treated with non-toxic concentration of **9** (VWK603) (0.63 and 1.25 µM) and the gene expressions of Bax and Bcl-2 were measured using quantitative RT-PCR. The Bax/Bcl-2 ratios of cells treated with 0.63 µM and 1.25 µM **9** (VWK603) compared to vehicle control were 1.54 and 1.59, respectively (Figure 3).

**Figure 3.**
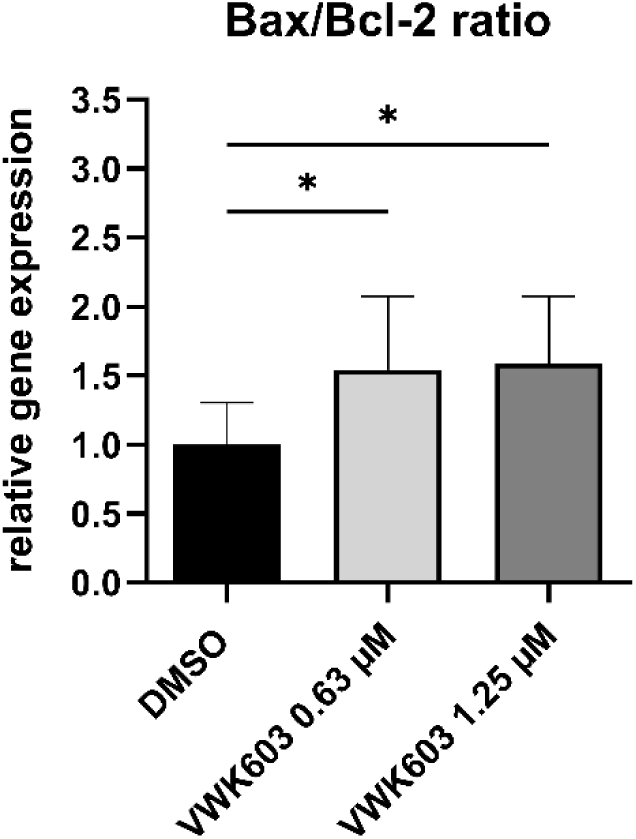
VWK603 impact on cellular Bax/Bcl-2 ratio in MDA-MB-231 cells. MDA-MB-231 cells were seeded and incubated for 24 hours before treatment. Cells were incubated with **9** (VWK603) at the displayed non-toxic concentrations or with DMSO for 72 hours. Subsequently, mRNA was isolated and transcribed into cDNA. Bax and Bcl-2 expression were measured by qRT-PCR using the 2^-ΔΔ^CT method and the Bax/Bcl-2 ratio was calculated, retrospectively. Statistical analysis was performed by using one-way ANOVA. P-values indicate the significant difference between with **9** (VWK603) and with DMSO treated cells. * = p ≤ 0.05

These data illustrate a significant increase of Bax/Bcl-2 ratio in MDA-MB-231-cells treated with **9** (VWK603) indicating an apoptosis induction of the novel drug via the mitochondrial way.

MDM2 inhibitors are known to cause on-target side effects in non-malignant cells.^5^ To evaluate the cytotoxicity of **9** (VWK603) in healthy tissue, we used the non-malignant breast epithelial cell line MCF-10A serving as a representative for non-malignant breast tissue. Cells were treated with different concentration of **9** (VWK603) or the reference compound Nutlin-3a. Both compounds reduced the cell viability in a dose-dependent manner in MCF-10A cells (Figure 4). However, in comparison to the studies performed in TNBC cells, the IC_50_ with 16.29 µM for **9** (VWK603) was essentially higher in non-malignant MCF-10A cells while for Nutlin-3a IC_50_ with 29.68 µM was more in the range of the TNBC cells. Thus, the ratio of IC_50_ of non-malignant breast to malignant TNBC cells was slightly higher for **9** (VWK603) (2.48 to 4.44) compared to Nutlin-3a (1.27 to 1.51). These data illustrate a lower toxicity of **9** (VWK603) in non-malignant vs. malignant cells with an even superior ratio for **9** (VWK603) compared to Nutlin-3a.

**Figure 4.**
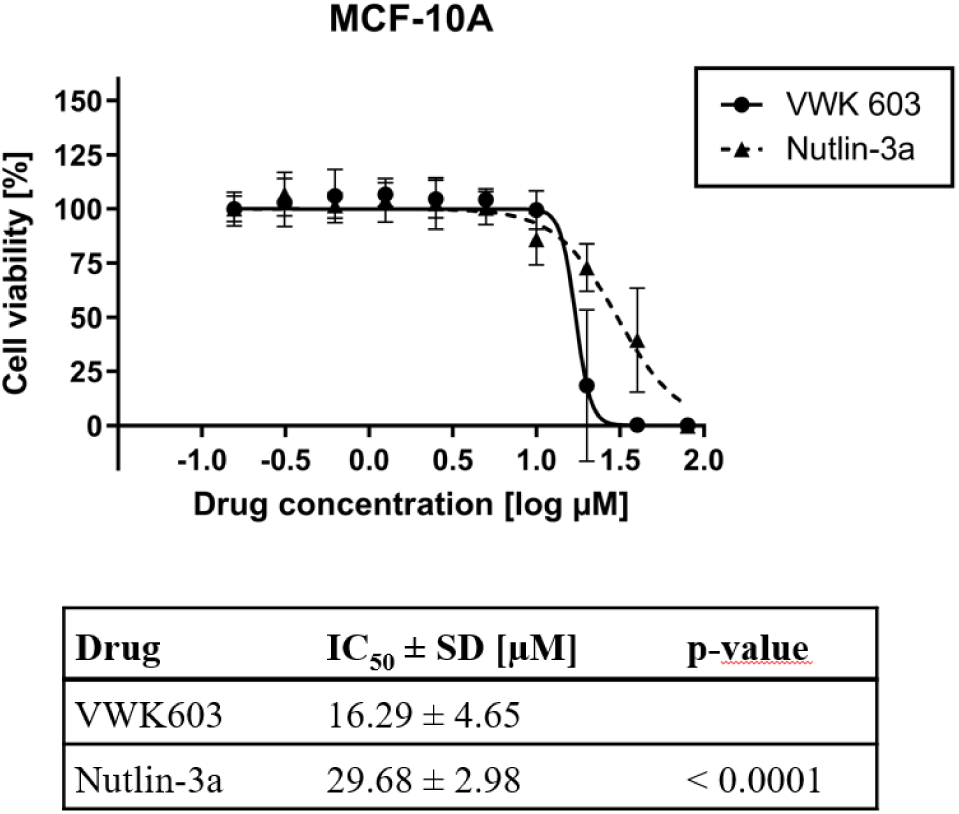
Toxicity studies of VWK603 in non-malignant MCF-10A cells. MCF-10A cells were seeded and incubated for 24 hours before treatment. **9** (VWK603) or Nutlin-3a was added at displayed concentrations and incubated for 72 hours. Treatment was repeated for another 72 hours. Subsequently, cell viability was determined by CellTiter Glo^®^ assay. The graph shows the mean ± SD n ≥ 3. Table below the graph contains the summarized IC_50_ values ± SD. Statistical analysis and significance were determined by using unpaired student’s t-test. P-value indicates the significant difference between the two compounds.

In this study, we identified the novel lead compound **9** (VWK603) to potently induce cell death in TNBC cell culture models. **9** has been designed to target MDM2 interactions with p53 and, due to the identical binding region^7, 8^, with p73 leading to activation of these tumor suppressors and induction of apoptosis. **9** (VWK603) induced cell death in different TNBC cell lines with an IC_50_ of 3.67 to 6.58 µM and showed significant superior activity compared to the MDM2 inhibitor Nutlin-3a (IC_50_: 19.36 to 23.15 µM). This anti-tumor effect was independent of functional p53 since the selected cell lines harbor p53-inactivating mutations and **9** (VWK603) equally induced cell death in p53^+/+^ as in p53^-/-^ cells. Since **9** (VWK603) significantly increased the Bax/Bcl-2 ratio, it occurs reasonable to expect apoptosis as the underlying mechanism of cell death observed in these studies like already widely described for other MDM2 inhibitors.^5,6^ Toxicity in non-malignant epithelial breast cells turned out to be considerably less (IC_50_: 16.29 µM) suggesting a possible tumor specificity of **9** (VWK603) and potential basis for further lead structure optimization towards a preclinical drug candidate.

The small molecule Nutlin-3a which is used in these studies as a reference compound is one of the most preclinically investigated MDM2 inhibitors which reactivates p53 in p53-wildtype cancers and like recently shown also p73 in p53-mutated cancer cells displaying significant antitumor activity^9, 20, 22, 23^ These observations are in line with reactivation of p73 reported to effectively induce apoptosis in TNBC cell lines^11, 32–35^ and compensation of mutation caused p53-functional absence by p73 upregulation^35^.

TNBC which mainly harbors inactivating p53 mutations (80%) is a highly aggressive breast cancer subtype^21^ where targeted therapeutic options are rare and not yet very efficient.^3^ This is mainly due to the fact that only few potential therapeutic targets have been discovered so far despite intensive basic research. Our studies suggest that the novel α-helix mimetic **9** (VWK603) represents a promising pharmacological lead compound to efficiently compromise TNBC cell viability by MDM2 targeting and has a significant potency for further preclinical drug development.

## Acknowledgements

The authors thank Ursula Grolik and Karin Juliane Bents for excellent technical support and Bert Vogelstein and Thomas G. Hofmann for supply of HCT116 p53^+/+^ and HCT116 p53^-/-^ cells.

## Statements & Declarations

### Author Contributions

Jasmin Linh On, Sanil Bhatia, Thomas Lenz, Andrea Kulik, Kai Stühler, Dieter Niederacher, Hans Neubauer, Tanja Fehm, Thomas Kurz and Knud Esser contributed to the study conception and design. Material preparation, data collection and analysis were performed by Jasmin Linh On, Vitalij Woloschin, Franziska Gier, Jia-Wey Tu, Sanil Bhatia. Synthesis of **9** (VWK603) was designed and performed by Vitalij Woloschin and Thomas Kurz. The first draft of the manuscript was written by Jasmin Linh On, Knud Esser and all authors commented on previous versions of the manuscript. All authors read and approved the final manuscript.

### Data availability

The datasets generated during and/or analysed during the current study are not publicly available due to a solely in-house electronic data archiving but are available from the corresponding author on reasonable request.

## Abbreviations

DMSO: dimethyl sulfoxide;
IC_50_: half-maximal inhibitory concentration;
MDM2: mouse double minute 2;
PARP: poly ADP ribose polymerase,
TNBC: triple-negative breast cancer

## Supporting information

### Materials and methods

All solvents and chemicals were used as purchased without further purification. The progress of all reactions was monitored on silica gel plates (with fluorescence indicator UV254) using the solven13t system stated. Flash column chromatography was carried out using CombiFlash® 200 (Teledyne Isco) and prepacked RediSep® (NP) silica cartridges. Melting points (Mp) were taken with a Büchi M-565 melting point apparatus in open capillaries and are uncorrected. ^1^H and ^13^C spectra were recorded on Avance™ DRX-500 (Bruker) (^1^H 500 MHz; ^13^C 126 MHz), Avance™ III–600 (Bruker) (^1^H 600 MHz; ^13^C 151 MHz) or Avance III – 300 (Bruker) (^1^H 300 MHz; ^13^C 75 MHz) spectrometers, respectively, using DMSO-*d*_6_ or Chlorodorm-*d* as solvents. Chemical shifts are given in parts million (ppm), (δ relative to residual solvent peak for ^1^H and ^13^C). ESI-MS was carried out using Bruker Daltonics UHR-QTOF maXis 4G (Bruker Daltonics) under electrospray ionization (ESI). Analytical HPLC was carried out on a Knauer HPLC system comprising of an Azura P6.1L pump, an Optimas 800 autosampler, a Fast Scanning Spectro-Photometer K-2600 and a Knauer Reversed Phase column (SN: FK36). Evaluated compounds were detected at 254 nm. The solvent gradient table is shown below. The purity of all final compounds was 95% or higher. Azlactone **1** was prepared as reported by MATOS *et al*. and crystallized from isopropyl alcohol^1^. Amidine hydrochloride **2** was prepared following a literature procedure by MARTINU *et al*.^2^ *Tert*-butyl 3-(2-hydroxyethyl)-1*H*-indole-1-carboxylate was synthesised using a procedure by GHOSH *et al*.^3^ **3** was synthesized according to a procedure previously described by us^4^.

### HPLC method

Prior to injection all compounds were dissolved either in acetonitrile or DMSO for a final concentration of 1 mg/mL. After the compound sample was injected onto the HPLC column through the autosampler an isocratic gradient of 90% water with 0.1% of TFA and 10% acetonitrile with 0.1% TFA was applied for 30 seconds. In the next step, a gradient was applied for 18.5 minutes where the water concentration was gradually reduced from 90% to 0%, while the acetonitrile concentration was increased from 10% to 100%. In the final phase an isocratic gradient of 100% acetonitrile with 0.1% of TFA was applied for 10 minutes. Flow rate: 1 mL/min. Detection: 254 nm.

### Cell cultures

All cell lines were obtained from ATCC (LGC Standards GmbH, Wesel, Deutschland). HCT116 p53^+/+^ and HCT116 p53^-/-^ were kindly provided by Dr. Bert Vogelstein (John Hopkins University, Baltimore, USA) and Dr. Thomas G. Hofmann (Institute of Toxicology, University Medical Center Mainz, Germany). MDA-MB-231 (ATCC®HTB-26™), MDA-MB-436 (ATCC®HTB-130™) MDA-MB-468 (ATCC®HTB-132™), NIH-3T3 (ATCC®CRL-1658™) were cultured with Roswell Park Memorial Institute (RPMI) Medium 1640 (Gibco, UK) supplemented with 10% fetal bovine serum (Gibco, USA) and 1% penicillin/streptomycin (Gibco, USA). HCT116 p53^+/+^ and HCT116 p53 ^-/-^ were cultured with McCoy 5a (Gibco, USA) with 2 mM L-Glutamine (Gibco, UK), 10% fetal bovine serum and 1% penicillin/streptomycin. MCF-10A cells (ATCC® CRL-10317™) were cultured with Dulbecco’s modified Eagle medium (DMEM) F12 (Gibco, USA) supplemented with 5% horse serum (Gibco, USA), 1% penicillin/streptomycin, epidermal growth factor (EGF) at 20 ng/mL (Miltenyi, Germany), hydrocortisone at 0.5 mg/mL (Sigma, #H-0888), cholera toxin at 100 ng/mL (Sigma, #C-8052) and insulin 10 µg/mL (Sigma, #I-1882). All cell lines were incubated in a humidified atmosphere at 37 °C in 5% CO2.

### Drugs and reagents

Nutlin-3a was purchased by Cayman, USA. VWK603 9 was synthesized by the authors T.K. and V.W. at the Institute of Pharmaceutical and Medicinal Chemistry, Heinrich-Heine-University Düsseldorf, Germany (detailed description in supplemental information). Nutlin-3a and VWK603 **9** were dissolved in dimethyl sulfoxide (Sigma, #D-2660) to achieve a final concentration of 10 mmol/L.

### Cell viability assay

Cells of all cell lines were seeded at a density of 2.6 x 10^4^ cells per well in 384 well microtiterplates (Greiner Bio-One, Austria) and incubated for 24 hours at 37 °C before treatment. After removing the culturing medium, serum-reduced medium RPMI 1640 advanced (Gibco, UK) with 2 mM L-Glutamine, phenol red (Sigma) and 100 nM dexamethasone (Sigma, #D-4902) was used for treatment. For MCF-10A cells, DMEM F12 supplemented with 2% horse serum, hydrocortisone at 0.5 mg/mL, cholera toxin at 100 ng/mL and insulin 10 µg/mL was used as assay medium. Compounds were added at mentioned concentrations and incubated for 72 hours and repeated for another 72 hours.

Subsequently, cell viability was evaluated using CellTiter Glo^®^ Luminescent Cell Viability Assay (Promega, USA) according to the manufacturer’s instructions. All measurements were performed using the Spark^®^ multimode microplate reader (Tecan, Switzerland).

### qRT-PCR

MDA-MB-231 cells were seeded at a density of 1.3 x 10^6^ cells per mL in a six well plate (Greiner Bio-One, Austria) and were incubated for 24 hours before treatment. After removing the culturing medium, a serum-free medium RPMI 1640 advanced with 2 mM L-Glutamine, phenol red and 100 nM dexamethasone was used for treatment. VWK603 9 was added at mentioned non-toxic concentrations and DMSO was used for control. Cells were incubated for 72 hours. Subsequently, mRNA was isolated using ReliaPrep™ RNA Cell Miniprep System (Promega, USA), and was transcribed into cDNA using Superscript^®^ First-Strand Synthesis System for RT-PCR (Thermo Fischer Scientific, USA) according to the manufacturer’s instructions. Primers obtained from Metabion (Germany) and LightCycler^®^ 480 SYBR® Green I Master (Roche, Germany) were used. qRT-PCR was performed using LightCycler^®^ 480 Instrument II (Roche, Germany). CT values were calculated according to the 2-^ΔΔ^ method. Sample values were normalized to the housekeeping genes GAPDH and HPRT.

### Statistical analysis

All experiments were performed at least three times. The results were expressed as the mean ± standard deviation (SD) of the mean. Student’s t-test or one-way ANOVA with Dunnett’s or Tukey’s post-hoc was performed. All analyses were conducted using GraphPad PRISM. A probability of error of p < 0.05 was considered statistically significant.

**Supplementary Table 1:**
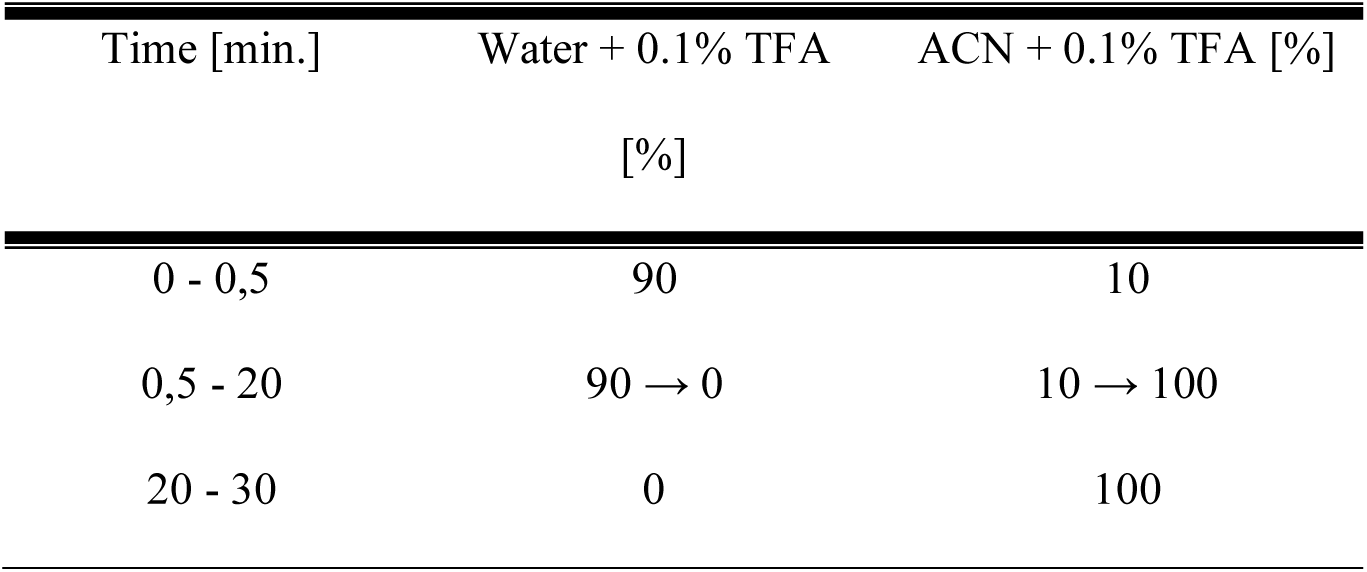
HPLC gradient.

#### General procedure for the synthesis of pyrimidines 4, 5

A suspension of 1.0 mmol **3**, 1.3 mmol triphenylphosphine and 1.3 mmol of the appropriate alcohol in 4 mL dry tetrahydrofuran was cooled to 0 °C using an ice bath. To the suspension 1.3 mmol of diisopropyl azodicarboxylate were added dropwise at 0 °C. After completion of the addition the ice bath was removed and the solution was stirred for 4 hours at room temperature. The solvent was removed under reduced pressure, the residue dissolved in 10 mL of diethyl ether and treated with 1 mL of n-hexane. The suspension was stirred for 30 minutes, the solid removed by filtering through a Büchner funnel and washed with 10 mL diethyl ether:n-hexane (1:1). The solvent was removed under reduced pressure and the residue purified using flash column chromatography (n-hexane:ethyl acetate 1:0→7:3→1:1).

#### Methyl 5-benzamido-4-(*sec*-butoxy)pyrimidine-2-carboxylate 4

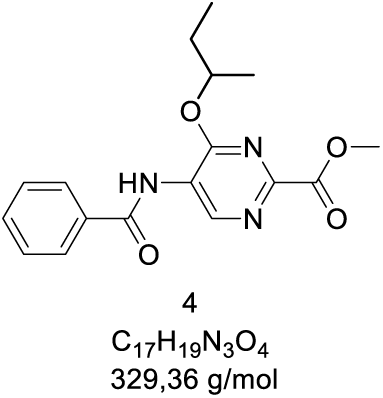

**Yield:** 45%; 5.92 g (18.0 mmol), colourless solid.

**Mp:** 98 °C (dichloromethane)

**HPLC:** R_t_: 12.77 min.; purity: 98.2%.

**^1^H NMR** (300 MHz, chloroform-*d*) δ 9.77 (s, 1H), 8.35 (s, 1H), 7.95 – 7.79 (m, 2H), 7.66 – 7.49 (m, 3H), 5.55 (h, J = 6.2 Hz, 1H), 4.01 (s, 3H), 1.94 – 1.73 (m, 2H), 1.43 (d, J = 6.1 Hz, 3H), 1.01 (t, J = 7.5 Hz, 3H).

**^13^C NMR** (75 MHz, chloroform-*d*) δ 165.48, 163.66, 158.45, 149.31, 144.75, 133.58, 132.85, 129.24, 127.23, 123.81, 76.53, 53.35, 28.95, 19.41, 9.72.

**Supplementary Fig. 1:**
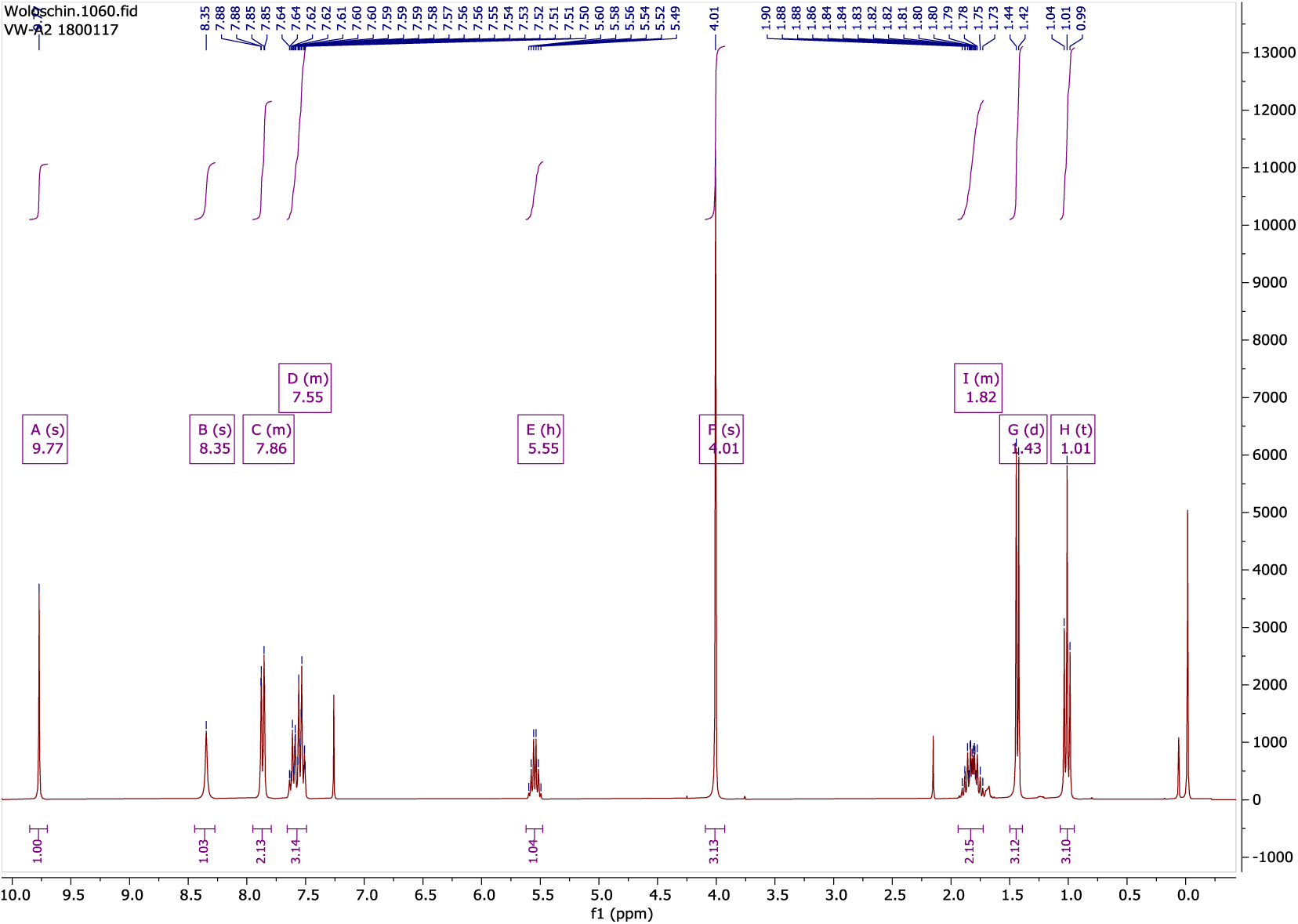
^1^H-NMR (300 MHz, chloroform-*d*) of **4** at room temperature.

**Supplementary Fig. 2:**
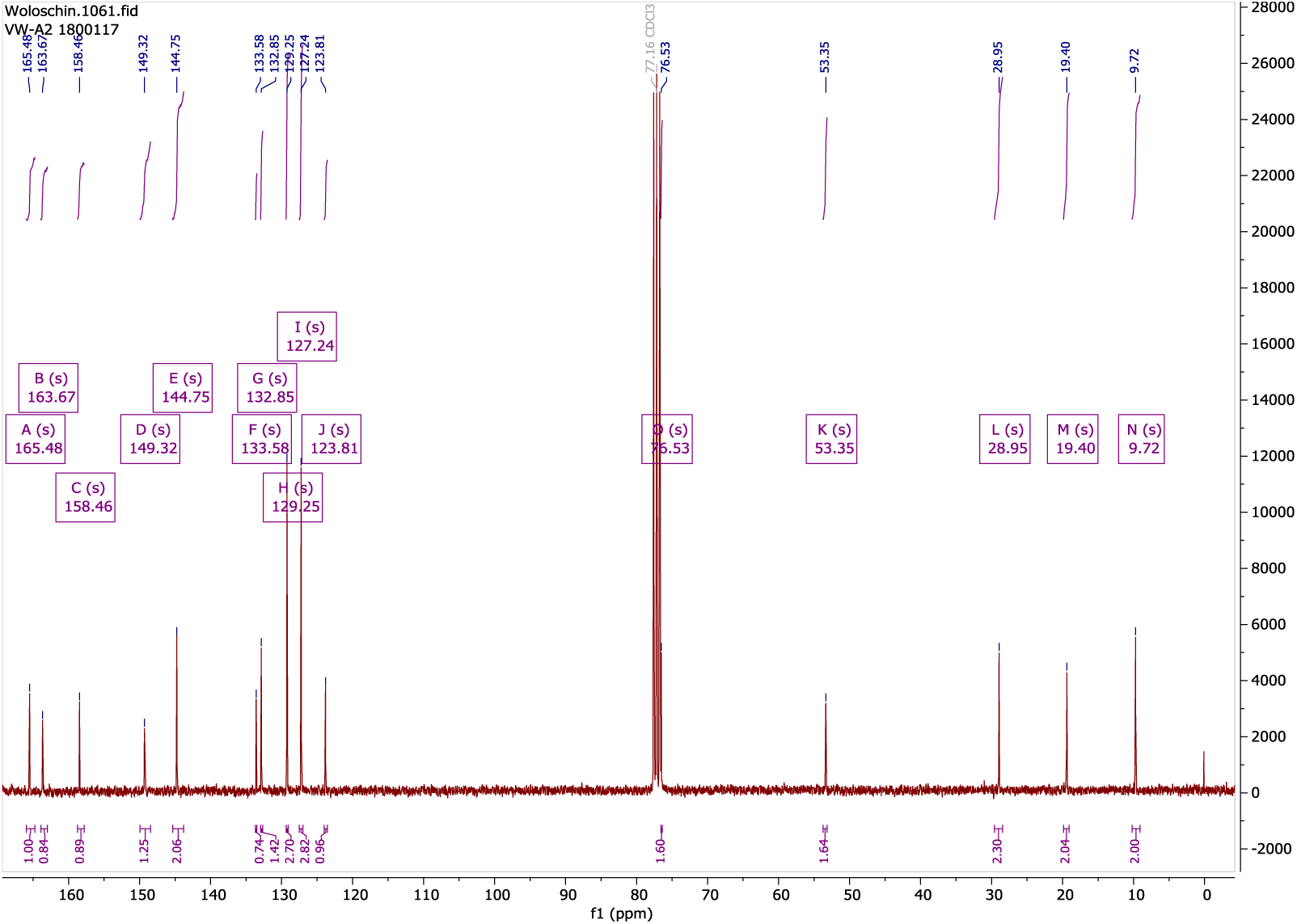
^13^C-NMR (75 MHz, chloroform-*d*) of 4 at room temperature.

*Tert*-butyl 3-(2-((5-benzamido-2-(methoxycarbonyl)pyrimidin-4-yl)oxy)ethyl)-1*H*-indole-1-carboxylate **5**

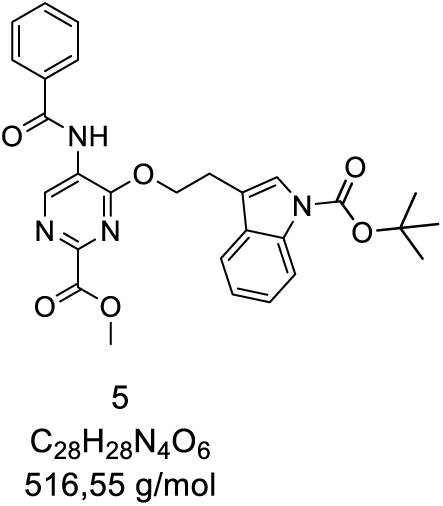

**Yield:** 65%, 1.47 g (2,85 mmol), yellow solid.

**Mp:** 158 °C (dichloromethane)

**HPLC:** R_t_: 18.05 min.; purity: 98.9%.

**^1^H NMR** (600 MHz, DMSO-*d*_6_) δ 9.92 (s, 1H), 9.16 (s, 1H), 8.02 (d, *J* = 8.2 Hz, 1H), 7.92 – 7.87 (m, 2H), 7.80 (d, *J* = 7.8 Hz, 1H), 7.65 (t, *J* = 7.4 Hz, 1H), 7.61 (s, 1H), 7.55 (t, *J* = 7.7 Hz, 2H), 7.29 (t, *J* = 7.7 Hz, 1H), 7.15 (t, *J* = 7.5 Hz, 1H), 4.75 (t, *J* = 6.8 Hz, 2H), 3.90 (s, 3H), 3.23 (t, *J* = 6.8 Hz, 2H), 1.57 (s, 9H).

**^13^C NMR** (75 MHz, chloroform-*d*) δ 165.51, 163.51, 158.39, 149.73, 149.20, 145.02, 135.53, 133.25, 132.81, 130.54, 129.22, 127.23, 124.83, 123.83, 123.59, 122.85, 118.89, 116.47, 115.54, 83.96, 67.39, 53.45, 28.30, 24.73.

**Supplementary Fig. 3:**
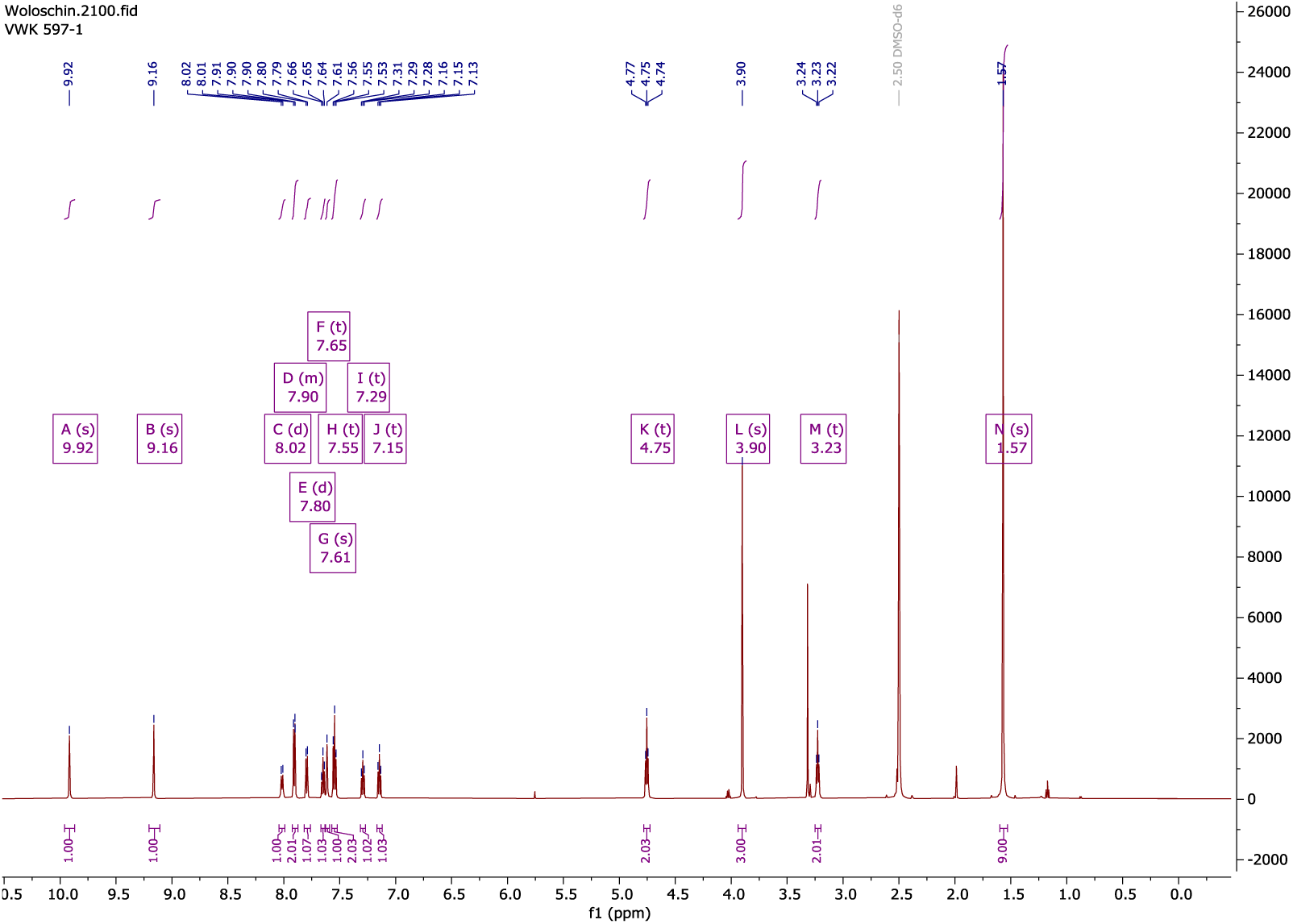
^1^H-NMR (600 MHz, DMSO-*d*_6_) of 5 at room temperature.

**Supplementary Fig. 4:**
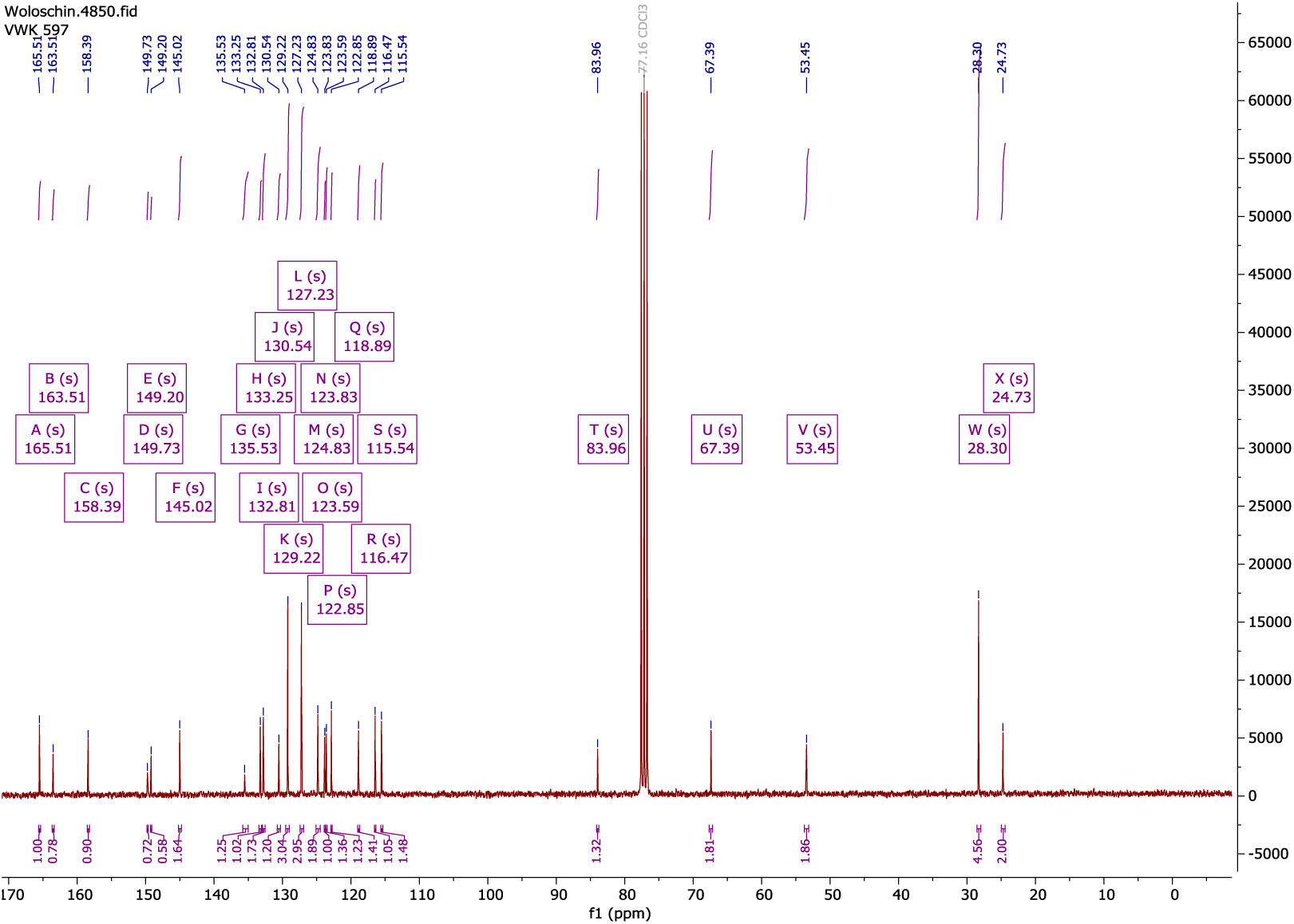
^13^C-NMR (75 MHz, chloroform-*d*) of 5 at room temperature.

#### Procedure for the synthesis of pyrimidine 6

To a solution of 6.59 g (20.0 mmol) of **4** in 100 mL of ethanol 14.1 g (160 mmol) of *N*,*N*-dimethylethylenediamine was added. The solution was stirred for 16 hours at room temperature, the solvent removed under reduced pressure and the residue dissolved in ethyl acetate (100 mL). The organic phase was washed with water (50 mL) and brine (50 mL) and dried with sodium sulfate. After filtration and removal of the solvent the product was finally purified using flash column chromatography (dichloromethane:MeOH).

#### 5-Benzamido-4-(*sec*-butoxy)-*N*-(2-(dimethylamino)ethyl)pyrimidine-2-carboxamide 6

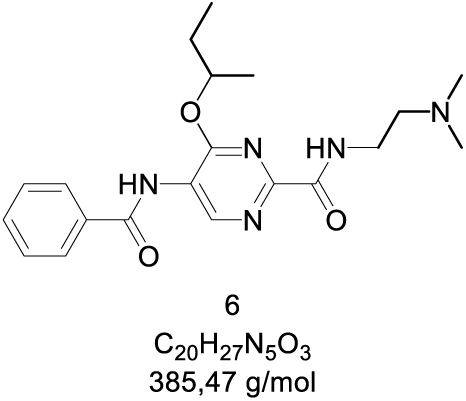

**Yield:** 79%; 6.12 g (15,9 mmol), colourless solid.

**Mp:** 125 °C (dichloromethane)

**HPLC:** R_t_: 8.92 min.; purity: 99.1%.

**^1^H NMR** (300 MHz, Acetone-*d*_6_) δ 9.53 (d, *J* = 2.1 Hz, 1H), 9.11 (s, 1H), 8.72 (t, *J* = 5.3 Hz, 1H), 8.05 – 7.90 (m, 2H), 7.72 – 7.50 (m, 3H), 5.41 (h, *J* = 6.2 Hz, 1H), 3.78 – 3.68 (m, 2H), 3.06 (t, *J* = 5.8 Hz, 2H), 2.71 (s, 6H), 1.75 (dtd, *J* = 6.8, 14.0, 21.6 Hz, 2H), 1.34 (d, *J* = 6.2 Hz, 3H), 0.92 (t, *J* = 7.4 Hz, 3H).

**^13^C NMR** (75 MHz, Acetone-*d*_6_) δ 166.55 (d, *J* = 6.1 Hz), 163.70 (d, *J* = 5.1 Hz), 160.21 (d, *J* = 4.8 Hz), 151.98, 146.67 (d, *J* = 8.8 Hz), 134.62 (d, *J* = 2.8 Hz), 133.40, 129.71, 128.49, 124.24 (d, *J* = 6.6 Hz), 77.05, 58.93, 45.12, 37.27 (d, *J* = 8.0 Hz), 29.29, 19.35, 9.90.

**Supplementary Figure 5:**
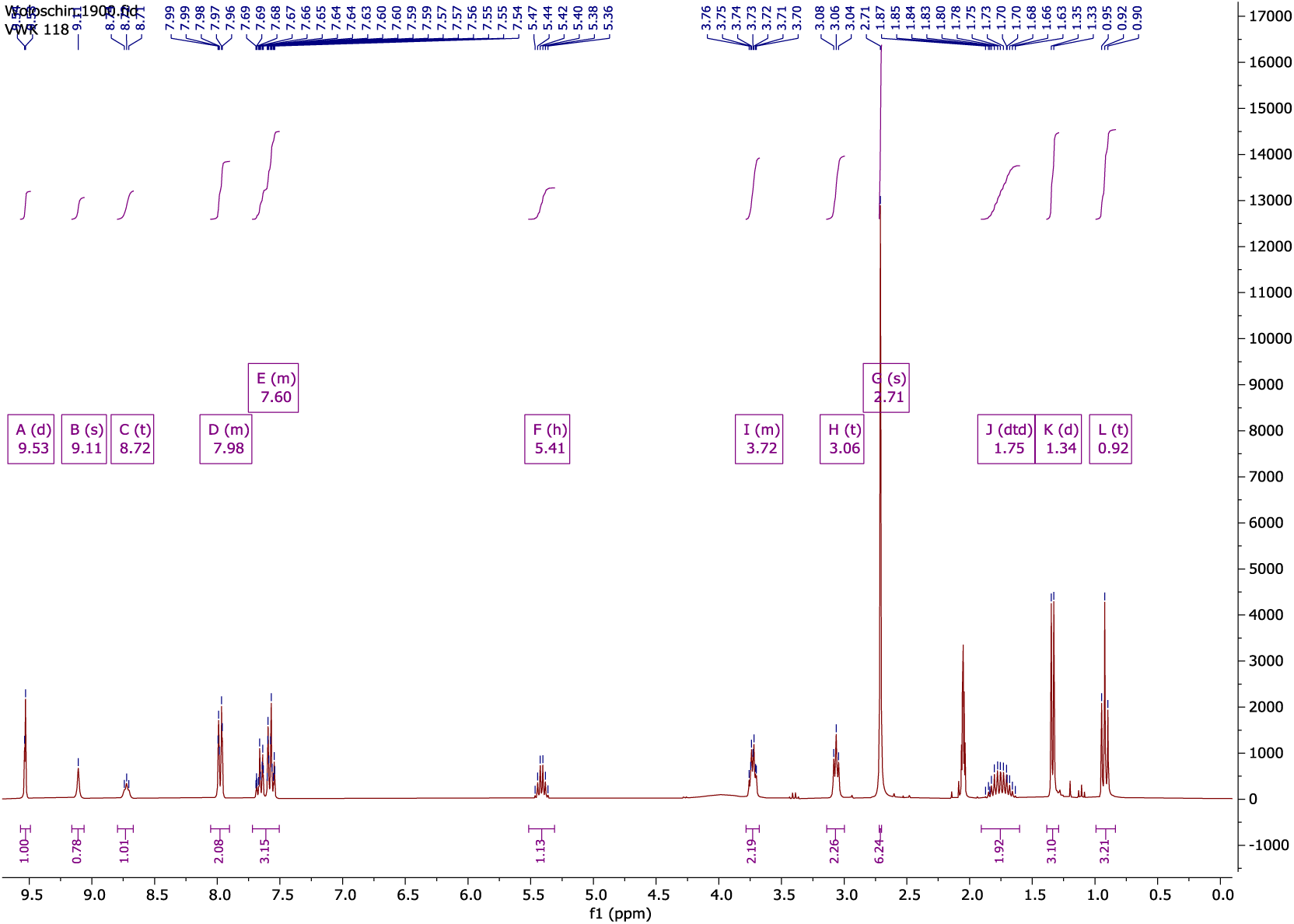
^1^H-NMR (300 MHz, acetone-*d*_6_) of **6** at room temperature.

**Supplementary Fig. 6:**
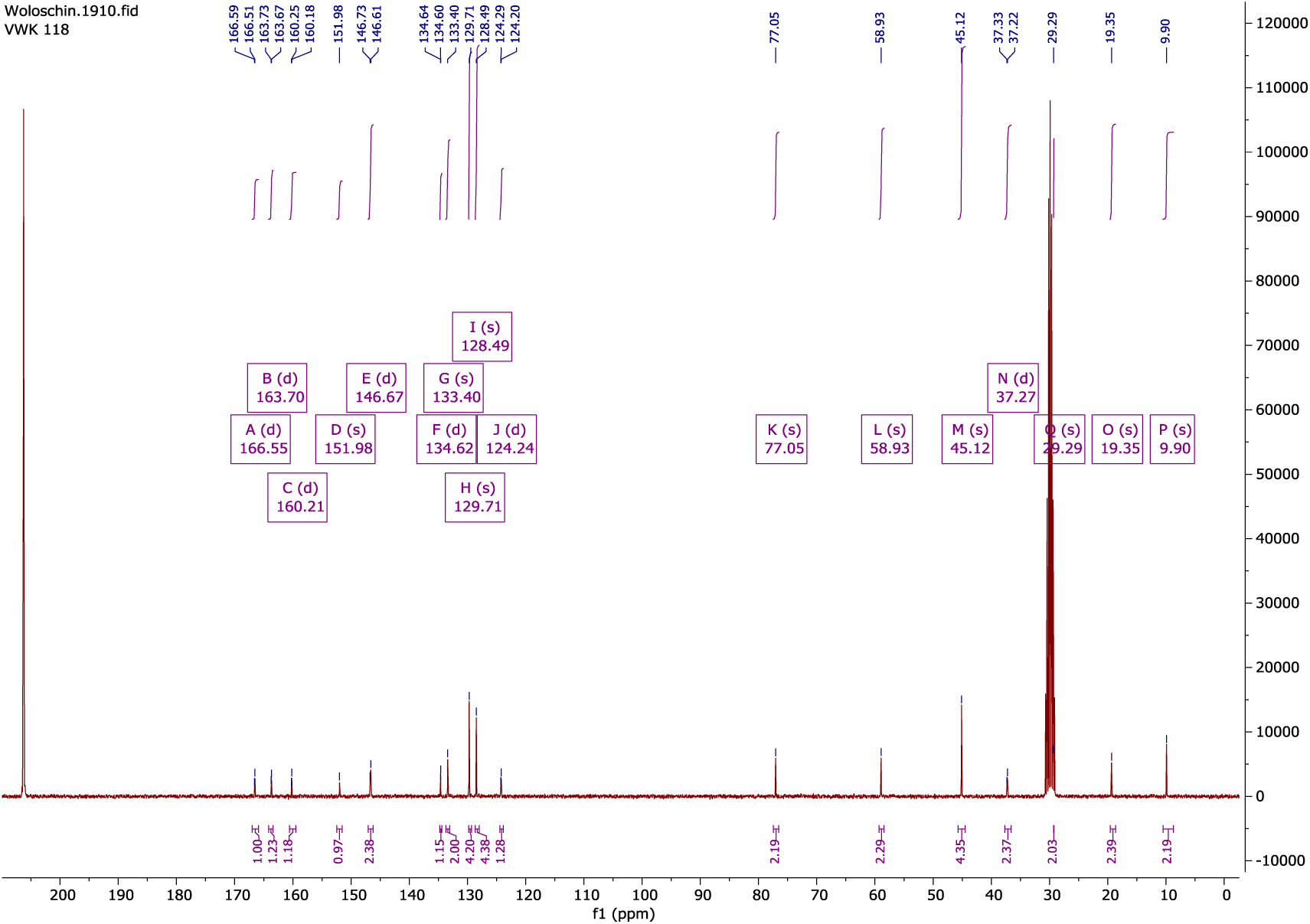
^13^C-NMR (75 MHz, acetone-*d*_6_) of 6 at room temperature.

#### Synthesis of 5-aminopyrimidine 7

To a solution of 1.94 g (5.03 mmol) **6** in 50 mL of dichloromethane, 0.26 g (6.54 mmol) of sodium hydroxide dissolved in 5 mL of methanol was added dropwise at room temperature. The solution was stirred for 24 hours at room temperature. The solvent was removed under reduced pressure und the residue suspended in a mixture of 20 mL ethyl acetate and 20 mL water. The mixture was stirred until all solids were dissolved. The aqueous layer was removed, the organic layer washed three times with a saturated sodium bicarbonate solution (10mL) and once with brine (10mL). The organic phase was dried with sodium sulfate and filtered. The solvent removed under reduced pressure and the solid finally recrystallized from n-hexane:2-propanol.

#### 5-Amino-4-(*sec*-butoxy)-*N*-(2-(dimethylamino)ethyl)pyrimidine-2-carboxamide 7

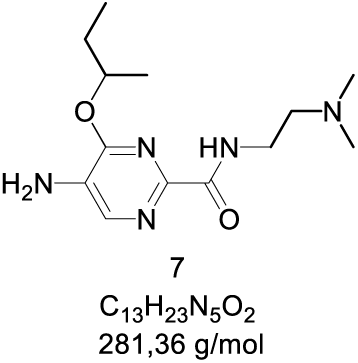

**Yield:** 81%; 1.15 g (4,09 mmol), colourless solid.

**Schmelzpunkt:** 170 °C (n-hexane:2-propanol)

**HPLC:** R_t_: 4.92 min.; purity: 99.9%.

**^1^H NMR** (300 MHz, chloroform-*d*) δ 8.12 (s, 1H), 7.97 (s, 1H), 5.34 (p, *J* = 6.2 Hz, 1H), 4.18 (s, 2H), 3.54 (q, *J* = 5.9 Hz, 2H), 2.55 (t, *J* = 6.1 Hz, 2H), 2.30 (s, 6H), 1.89 – 1.59 (m, 2H), 1.36 (d, *J* = 6.2 Hz, 3H), 0.96 (t, *J* = 7.5 Hz, 3H).

**^13^C NMR** (75 MHz, chloroform-*d*) δ 163.15, 156.72, 146.61, 137.95, 131.09, 74.72, 58.02, 45.30, 37.11, 28.93, 19.37, 9.71.

**Supplementary Fig. 7:**
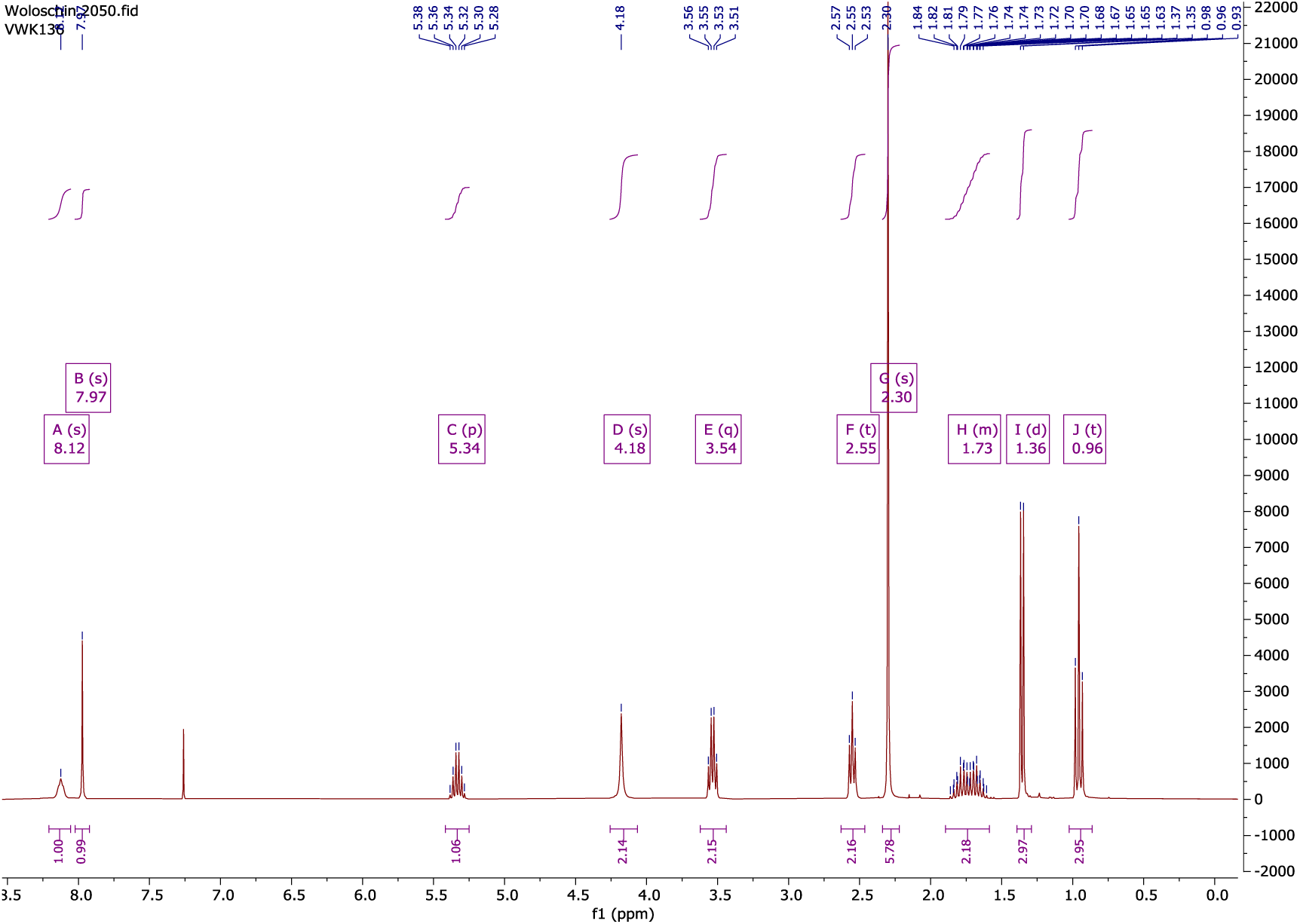
^1^H-NMR (300 MHz, chloroform-*d*) of 7 at room temperature.

**Supplementary Fig.8:**
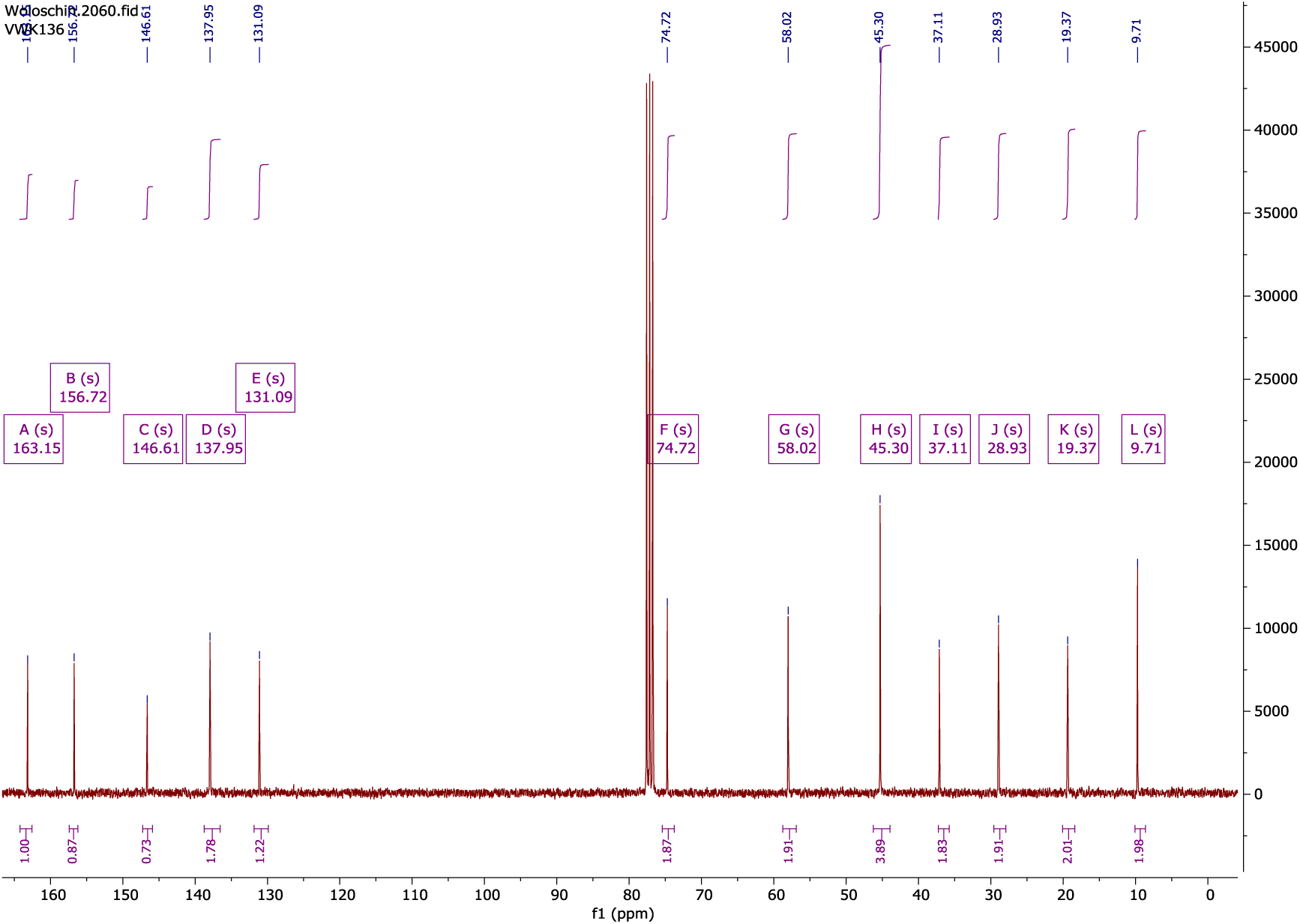
^13^C-NMR (75 MHz, chloroform-*d*) of 7 at room temperature.

#### Synthesis of pyrimidine 8

To a solution of 517 mg (1.00 mmol) methyl ester **5** in 5 mL of tetrahydrofuran was added 1 mL of a 1 M solution of lithium hydroxide in water. The solution was stirred for 24 hours at room temperature. The solvent was removed, the residue suspended in 30 mL diethyl ether and submerged for 5 min. in an ultra-sonic bath. Finally, the solid was filtered, washed with 10 mL of diethyl ether and dried under vacuum.

Lithium 5-benzamido-4-(2-(1-(*tert*-butoxycarbonyl)-1*H*-indol-3-yl)ethoxy)pyrimidine-2-carboxylate **8**

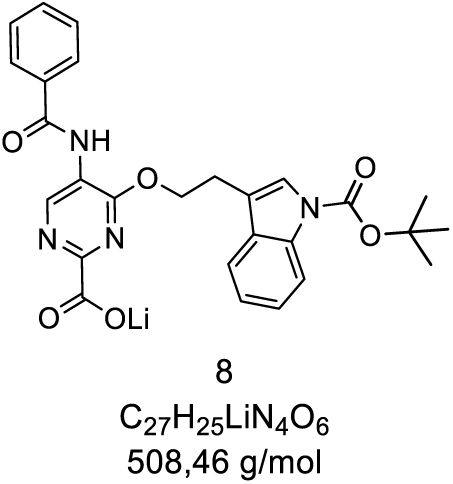

**Yield:** 86%; 438 mg (0.861 mmol), colourless solid.

**Mp:** 206 °C (diethyl ether)

**HPLC:** R_t_: 15.57 min.; purity: 99.5%.

**^1^H NMR** (600 MHz, DMSO-*d*_6_) δ 9.83 (s, 1H), 8.79 (s, 1H), 8.01 (d, *J* = 8.3 Hz, 1H), 7.93 (d, *J* = 7.9 Hz, 2H), 7.80 (d, *J* = 7.8 Hz, 1H), 7.65 – 7.56 (m, 2H), 7.51 (t, *J* = 7.6 Hz, 2H), 7.27 (t, *J* = 7.7 Hz, 1H), 7.10 (t, *J* = 7.5 Hz, 1H), 4.69 (t, *J* = 6.7 Hz, 2H), 3.18 (t, *J* = 6.7 Hz, 2H), 1.56 (s, 9H).

**^13^C NMR** (75 MHz, DMSO-*d*_6_) δ 166.25, 165.59, 161.59, 149.90, 149.01, 134.63, 133.81, 131.86, 130.25, 128.42, 127.75, 124.31, 123.48, 122.52, 119.65, 117.06, 114.58, 83.45, 79.19, 27.64, 23.90.

**Supplementary Fig. 9:**
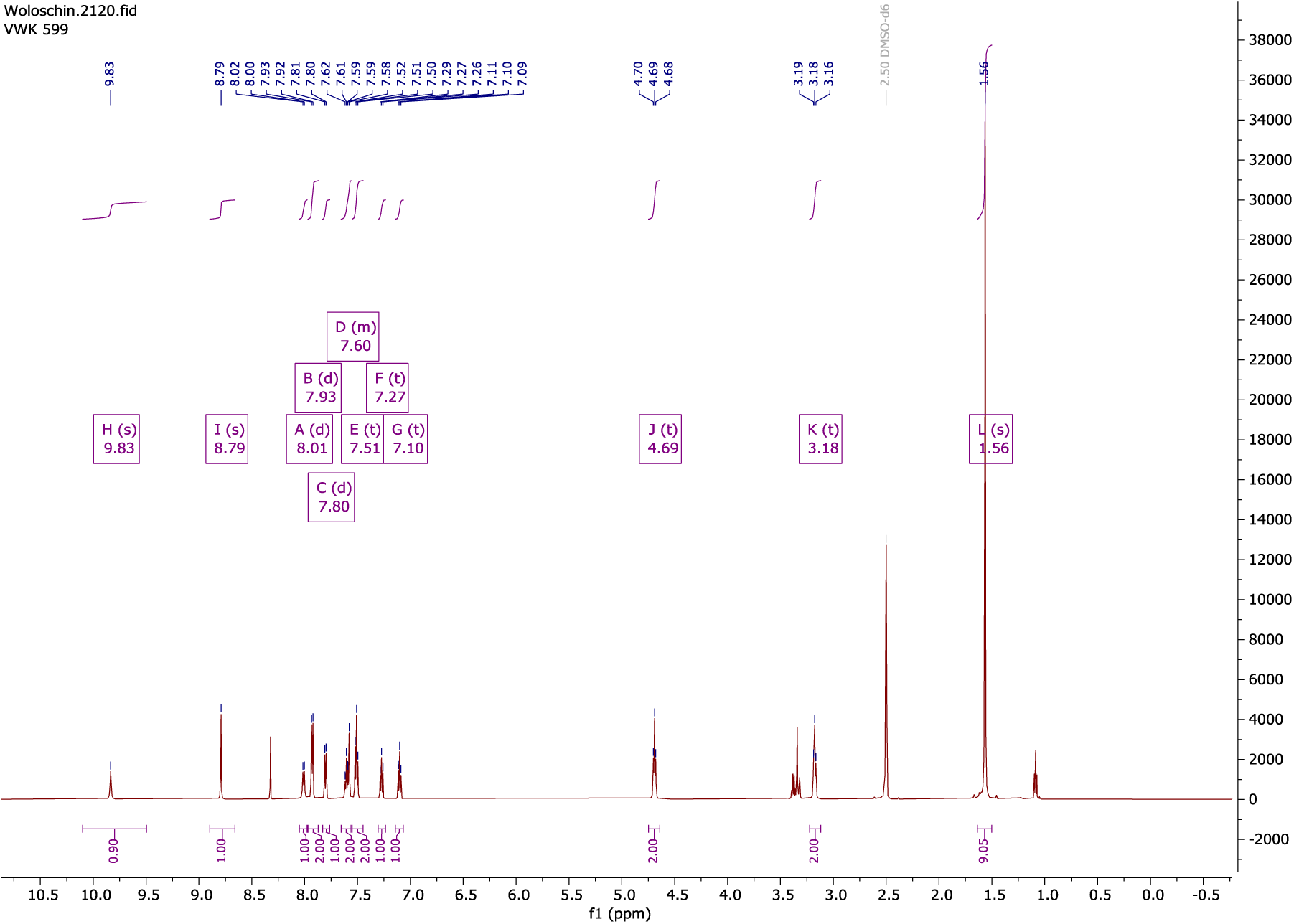
^1^H-NMR (600 MHz, DMSO-*d*_6_) of 8 at room temperature.

**Supplementary Fig. 10:**
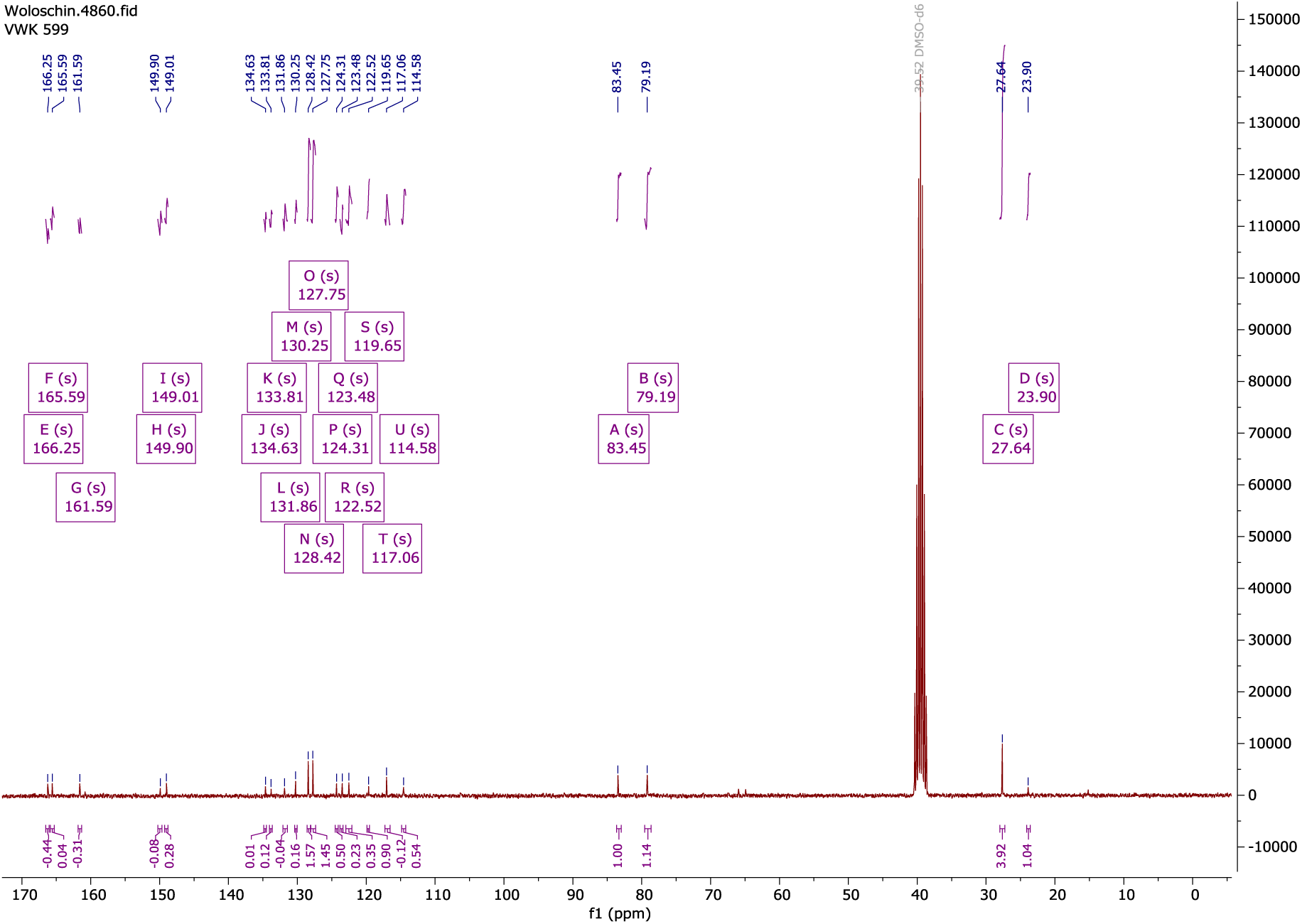
^13^C-NMR (75 MHz, DMSO-*d*_6_) of 8 at room temperature.

#### Synthesis of bipyrimidineamide 9

211 mg (0.75 mmol) **7**, 419 mg (0.83 mmol) **8** and 316 mg (0.83 mmol) HATU were dissolved in 3 mL of dimethylformamide. The solution was stirred for 24 hours and then diluted with 20 mL of ethyl acetate. The organic phase was washed once with 10 mL of a saturated sodium carbonate solution, 10 mL of brine, dried with sodium sulfate and filtered. After removing the solvent under reduced pressure, the product was purified using flash column chromatography (dichloromethane:MeOH).

*Tert*-butyl 3-(2-((5-benzamido-2-((4-(*sec*-butoxy)-2-((2-(dimethylamino)ethyl)carbamoyl)pyrimidin-5-yl)carbamoyl)pyrimidin-4-yl)oxy)ethyl)-1*H*-indole-1-carboxylate **9**

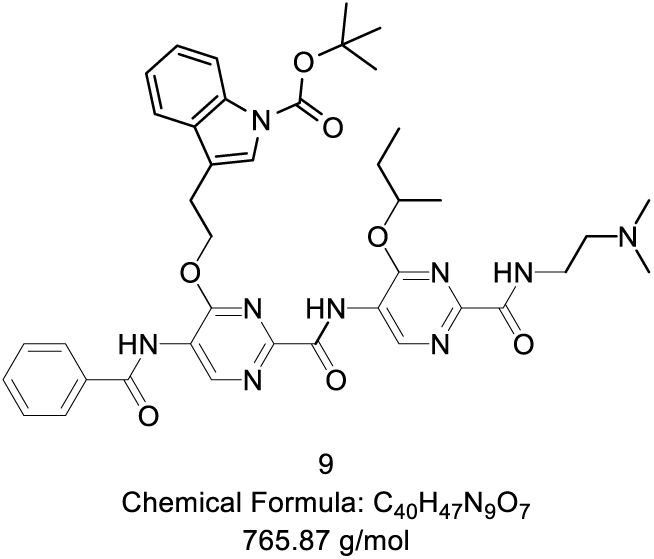

**Yield:** 29%; 165 mg (0.215 mmol), yellow solid.

**Mp:** 129 °C (dichloromethane)

**HPLC:** R_t_: 14.75 min.; purity 97.4%.

**^1^H NMR** (600 MHz, chloroform-*d*) δ 11.57 (s, 1H), 10.31 (s, 1H), 9.79 (s, 1H), 9.78 (s, 1H), 8.87 (t, *J* = 6.2 Hz, 1H), 8.22 (s, 1H), 8.13 (s, 1H), 7.68 (d, *J* = 7.6 Hz, 2H), 7.60 (t, *J* = 8.2 Hz, 2H), 7.55 (s, 1H), 7.50 (t, *J* = 7.7 Hz, 2H), 7.31 (t, *J* = 7.7 Hz, 1H), 7.18 (t, *J* = 7.5 Hz, 1H), 5.54 (h, *J* = 6.2 Hz, 1H), 4.99 (q, *J* = 6.4 Hz, 2H), 3.98 – 3.89 (m, 2H), 3.46 (t, *J* = 5.3 Hz, 2H), 3.32 (t, *J* = 6.5 Hz, 2H), 2.98 (s, 6H), 1.79 – 1.69 (m, 2H), 1.63 (s, 9H), 1.37 (d, *J* = 6.2 Hz, 3H), 0.98 (t, *J* = 7.4 Hz, 3H).

**^13^C NMR** (75 MHz, chloroform-*d*) δ 165.62, 163.94, 161.37, 160.87, 160.19, 158.83, 158.76, 149.85, 149.74, 149.56, 144.55, 143.79, 135.53, 133.09, 132.95, 130.47, 129.29, 127.26, 124.93, 123.90, 123.68, 123.40, 122.93, 118.73, 116.34, 115.61, 84.09, 77.36, 67.69, 57.58, 44.08, 35.57, 28.81, 28.28, 24.71, 19.08, 9.32.

**HR-MS (ESI+):** calculated for [C_40_H_47_N_9_O_7_+H^+^] m/z: 766.3671; found: 766.3679

**Supplementary Fig. 11:**
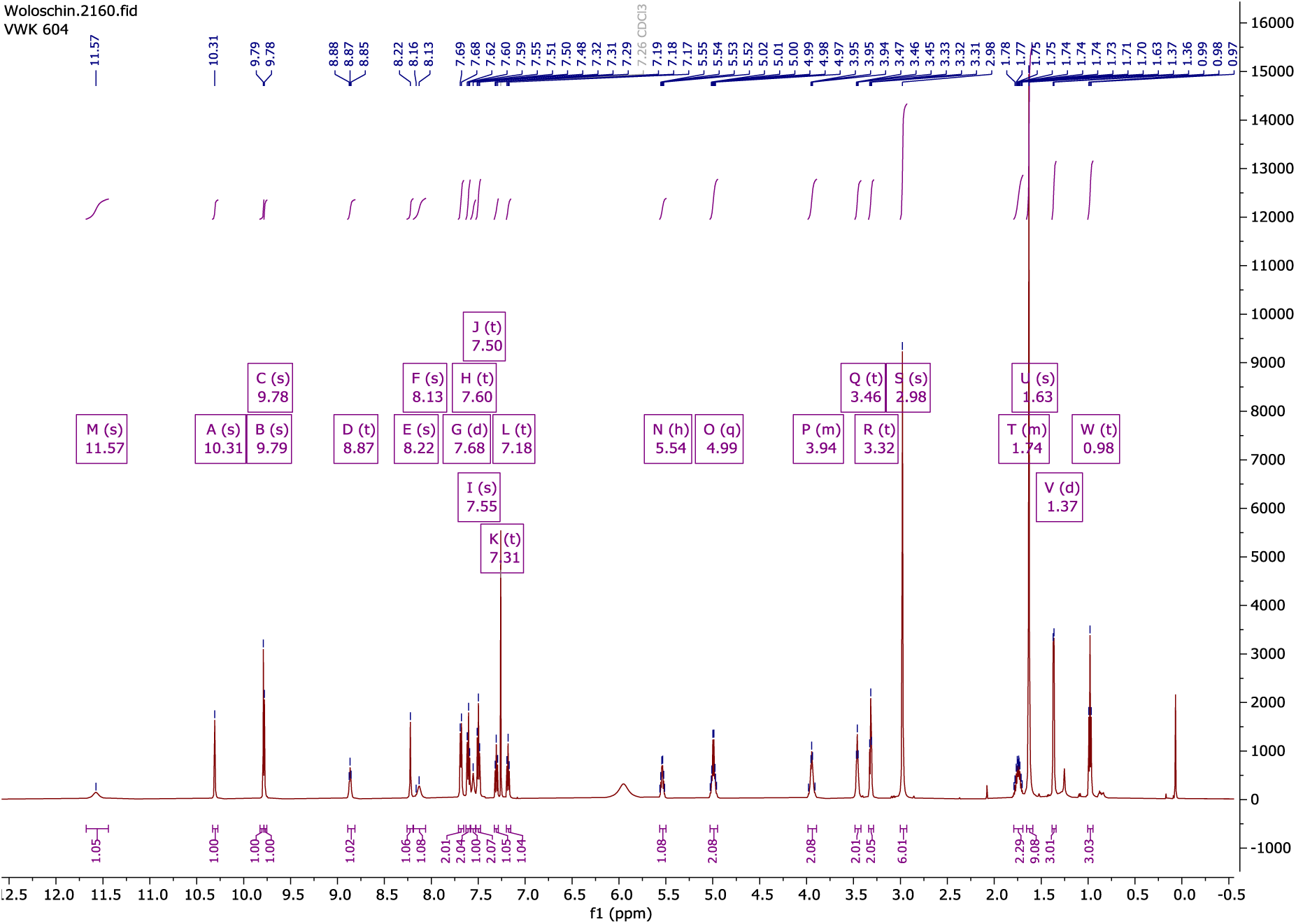
^1^H-NMR (600 MHz, chloroform-*d*) of 9 at room temperature.

**Supplementary Fig. 12:**
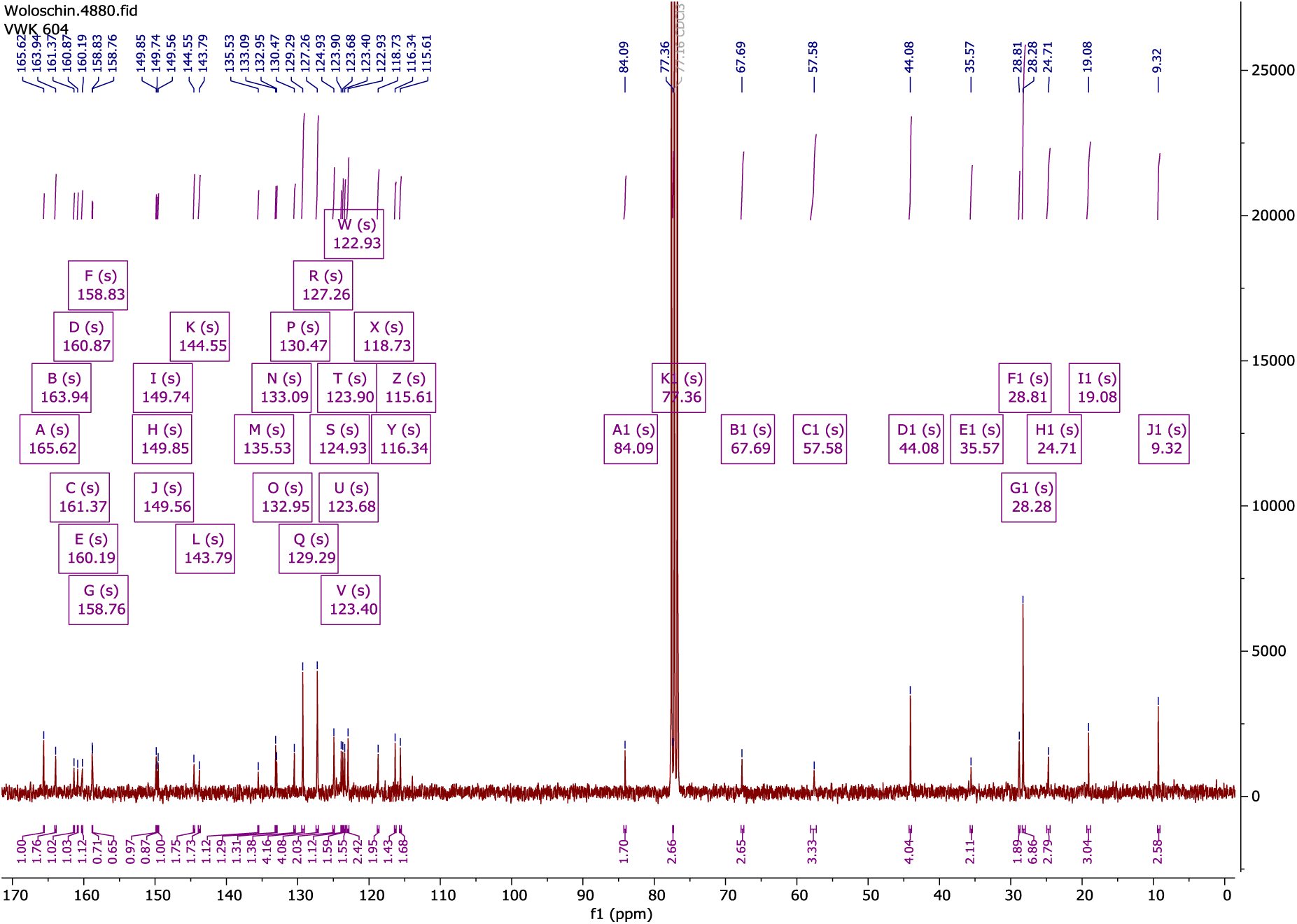
^13^C-NMR (75 MHz, chloroform-*d*) of 9 at room temperature.

**Supplementary Fig. 13:**
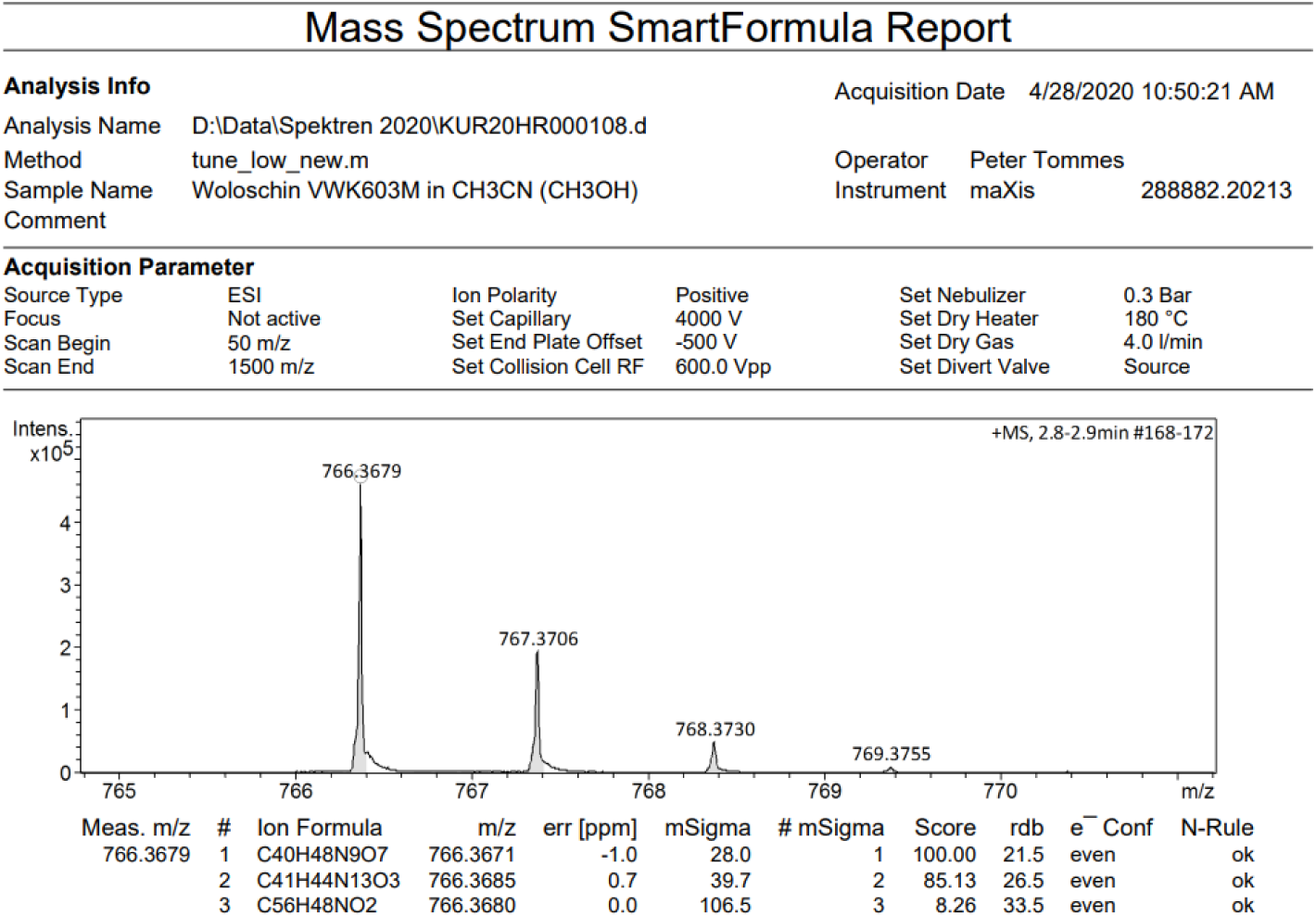
ESI spectrometry measurement of 9.

## References

(1) Yin, L.; Duan, J. J.; Bian, X. W.; Yu, S. C. Triple-negative breast cancer molecular subtyping and treatment progress. Breast Cancer Res 2020, 22 (1), 61.

(2) Ribeiro, R.; Carvalho, M. J.; Goncalves, J.; Moreira, J. N. Immunotherapy in triple-negative breast cancer: Insights into tumor immune landscape and therapeutic opportunities. Front Mol Biosci 2022, 9, 903065.

(3) Li, Y.; Zhang, H.; Merkher, Y.; Chen, L.; Liu, N.; Leonov, S.; Chen, Y. Recent advances in therapeutic strategies for triple-negative breast cancer. J. Hematol. Oncol. 2022, 15 (1), 121.

(4) Nag, S.; Qin, J.; Srivenugopal, K. S.; Wang, M.; Zhang, R. The MDM2-p53 pathway revisited. J Biomed Res 2013, 27 (4), 254–271.

(5) Konopleva, M.; Martinelli, G.; Daver, N.; Papayannidis, C.; Wei, A.; Higgins, B.; Ott, M.; Mascarenhas, J.; Andreeff, M. MDM2 inhibition: an important step forward in cancer therapy. Leukemia 2020, 34 (11), 2858–2874.

(6) Wang, S.; Chen, F. E. Small-molecule MDM2 inhibitors in clinical trials for cancer therapy. Eur. J. Med. Chem. 2022, 236, 114334.

(7) Bálint, E.; Bates, S.; Vousden, K. H. Mdm2 binds p73 alpha without targeting degradation. Oncogene 1999, 18 (27), 3923–3929.

(8) Dobbelstein, M.; Wienzek, S.; König, C.; Roth, J. Inactivation of the p53-homologue p73 by the mdm2-oncoprotein. Oncogene 1999, 18 (12), 2101–2106.

(9) Lau, L. M.; Nugent, J. K.; Zhao, X.; Irwin, M. S. HDM2 antagonist Nutlin-3 disrupts p73-HDM2 binding and enhances p73 function. Oncogene 2008, 27 (7), 997–1003.

(10) Peirce, S. K.; Findley, H. W. The MDM2 antagonist nutlin-3 sensitizes p53-null neuroblastoma cells to doxorubicin via E2F1 and TAp73. Int J Oncol 2009, 34 (5), 1395–1402.

(11) Adams, C. M.; Mitra, R.; Xiao, Y.; Michener, P.; Palazzo, J.; Chao, A.; Gour, J.; Cassel, J.; Salvino, J. M.; Eischen, C. M. Targeted MDM2 Degradation Reveals a New Vulnerability for p53-Inactivated Triple-Negative Breast Cancer. Cancer Discov 2023, 13 (5), 1210–1229.

(12) Pelay-Gimeno, M.; Glas, A.; Koch, O.; Grossmann, T. N. Structure-Based Design of Inhibitors of Protein-Protein Interactions: Mimicking Peptide Binding Epitopes. Angew. Chem. Int. Ed. Engl. 2015, 54 (31), 8896–8927.

(13) Bullock, B. N.; Jochim, A. L.; Arora, P. S. Assessing Helical Protein Interfaces for Inhibitor Design. J. Am. Chem. Soc. 2011, 133 (36), 14220–14223.

(14) Mabonga, L.; Kappo, A. P. Peptidomimetics: A Synthetic Tool for Inhibiting Protein–Protein Interactions in Cancer. Int. J. Pept. Res. Ther. 2019, 26 (1), 225–241.

(15) Lanning, M.; Fletcher, S. Recapitulating the alpha-helix: nonpeptidic, low-molecular-weight ligands for the modulation of helix-mediated protein-protein interactions. Future Med. Chem. 2013, 5 (18), 2157–2174.

(16) Saraogi, I.; Hamilton, A. D. alpha-Helix mimetics as inhibitors of protein-protein interactions. Biochem Soc Trans 2008, 36 (Pt 6), 1414–1417.

(17) Kussie, P. H.; Gorina, S.; Marechal, V.; Elenbaas, B.; Moreau, J.; Levine, A. J.; Pavletich, N. P. Structure of the MDM2 oncoprotein bound to the p53 tumor suppressor transactivation domain. Science 1996, 274 (5289), 948–953.

(18) Bhatia, S.; Spanier, L.; Bickel, D.; Dienstbier, N.; Woloschin, V.; Vogt, M.; Pols, H.; Lungerich, B.; Reiners, J.; Aghaallaei, N.;, et al. Development of a First-in-Class Small-Molecule Inhibitor of the C-Terminal Hsp90 Dimerization. ACS Cent Sci 2022, 8 (5), 636–655.

(19) Spanier, L.; Ciglia, E.; Hansen, F. K.; Kuna, K.; Frank, W.; Gohlke, H.; Kurz, T. Design, synthesis, and conformational analysis of trispyrimidonamides as α-helix mimetics. J. Org. Chem. 2014, 79 (4), 1582–1593.

(20) Vassilev, L. T.; Vu, B. T.; Graves, B.; Carvajal, D.; Podlaski, F.; Filipovic, Z.; Kong, N.; Kammlott, U.; Lukacs, C.; Klein, C.;, et al. In vivo activation of the p53 pathway by small-molecule antagonists of MDM2. Science 2004, 303 (5659), 844–848.

(21) Cancer Genome Atlas, N. Comprehensive molecular portraits of human breast tumours. Nature 2012, 490 (7418), 61–70.

(22) Wang, J.; Zheng, T.; Chen, X.; Song, X.; Meng, X.; Bhatta, N.; Pan, S.; Jiang, H.; Liu, L. MDM2 antagonist can inhibit tumor growth in hepatocellular carcinoma with different types of p53 in vitro. J. Gastroenterol. Hepatol. 2011, 26 (2), 371–377.

(23) Tonsing-Carter, E.; Bailey, B. J.; Saadatzadeh, M. R.; Ding, J.; Wang, H.; Sinn, A. L.; Peterman, K. M.; Spragins, T. K.; Silver, J. M.; Sprouse, A. A.;, et al. Potentiation of Carboplatin-Mediated DNA Damage by the Mdm2 Modulator Nutlin-3a in a Humanized Orthotopic Breast-to-Lung Metastatic Model. Mol Cancer Ther 2015, 14 (12), 2850–2863.

(24) Gomes, S.; Raimundo, L.; Soares, J.; Loureiro, J. B.; Leao, M.; Ramos, H.; Monteiro, M. N.; Lemos, A.; Moreira, J.; Pinto, M.;, et al. New inhibitor of the TAp73 interaction with MDM2 and mutant p53 with promising antitumor activity against neuroblastoma. Cancer Lett 2019, 446, 90–102.

(25) Wang, W.; Qin, J. J.; Voruganti, S.; Srivenugopal, K. S.; Nag, S.; Patil, S.; Sharma, H.; Wang, M. H.; Wang, H.; Buolamwini, J. K.;, et al. The pyrido[b]indole MDM2 inhibitor SP-141 exerts potent therapeutic effects in breast cancer models. Nat Commun 2014, 5, 5086.

(26) Fan, Y.; Li, M.; Ma, K.; Hu, Y.; Jing, J.; Shi, Y.; Li, E.; Dong, D. Dual-target MDM2/MDMX inhibitor increases the sensitization of doxorubicin and inhibits migration and invasion abilities of triple-negative breast cancer cells through activation of TAB1/TAK1/p38 MAPK pathway. Cancer Biol Ther 2019, 20 (5), 617–632.

(27) Fan, S.; Cherney, B.; Reinhold, W.; Rucker, K.; O’Connor, P. M. Disruption of p53 function in immortalized human cells does not affect survival or apoptosis after taxol or vincristine treatment. Clin Cancer Res 1998, 4 (4), 1047–1054.

(28) Dudgeon, D. D.; Shinde, S.; Hua, Y.; Shun, T. Y.; Lazo, J. S.; Strock, C. J.; Giuliano, K. A.; Taylor, D. L.; Johnston, P. A.; Johnston, P. A. Implementation of a 220,000-compound HCS campaign to identify disruptors of the interaction between p53 and hDM2 and characterization of the confirmed hits. J. Biomol. Screen. 2010, 15 (7), 766–782.

(29) Oltvai, Z. N.; Milliman, C. L.; Korsmeyer, S. J. Bcl-2 heterodimerizes in vivo with a conserved homolog, Bax, that accelerates programmed cell death. Cell 1993, 74 (4), 609–619.

(30) Akhtar, S.; Khan, M. K. A.; Arif, J. M. Evaluation and Elucidation Studies of Natural Aglycones for Anticancer Potential using Apoptosis-Related Markers: An In silico Study. Interdiscip Sci 2018, 10 (2), 297–310.

(31) Raisova, M.; Hossini, A. M.; Eberle, J.; Riebeling, C.; Wieder, T.; Sturm, I.; Daniel, P. T.; Orfanos, C. E.; Geilen, C. C. The Bax/Bcl-2 ratio determines the susceptibility of human melanoma cells to CD95/Fas-mediated apoptosis. J. Invest. Dermatol. 2001, 117 (2), 333–340.

(32) Tiwary, R.; Yu, W.; Sanders, B. G.; Kline, K. α-TEA cooperates with chemotherapeutic agents to induce apoptosis of p53 mutant, triple-negative human breast cancer cells via activating p73. Breast Cancer Res 2011, 13 (1), R1.

(33) Choi, E. K.; Kim, S. M.; Hong, S. W.; Moon, J. H.; Shin, J. S.; Kim, J. H.; Hwang, I. Y.; Jung, S. A.; Lee, D. H.; Lee, E. Y.;, et al. SH003 selectively induces p73dependent apoptosis in triplenegative breast cancer cells. Mol Med Rep 2016, 14 (4), 3955–3960.

(34) Huang, L.; Li, A.; Liao, G.; Yang, F.; Yang, J.; Chen, X.; Jiang, X. Curcumol triggers apoptosis of p53 mutant triple-negative human breast cancer MDA-MB 231 cells via activation of p73 and PUMA. Oncol Lett 2017, 14 (1), 1080–1088.

(35) Vayssade, M.; Haddada, H.; Faridoni-Laurens, L.; Tourpin, S.; Valent, A.; Benard, J.; Ahomadegbe, J. C. P73 functionally replaces p53 in Adriamycin-treated, p53-deficient breast cancer cells. Int J Cancer 2005, 116 (6), 860–869.

## References

(1) Norton Matos, M. R. P.; Gois, P. M. P.; Mata, M. L. E. N.; Cabrita, E. J.; Afonso, C. A.M. Studies on the Preparation of 4-Ethoxyalkyliden and 4-Aminoalkyliden-5(4H)-oxazolones. Synthetic Communications 2003, 33 (8), 1285–1299.

(2) Martinu, T.; Dailey, W. P. Synthesis of carboalkoxychloro- and bromodiazirines. J. Org. Chem. 2004, 69 (21), 7359–7362.

(3) Ghosh, N.; Nayak, S.; Sahoo, A. K. Gold-catalyzed regioselective hydration of propargyl acetates assisted by a neighboring carbonyl group: access to α-acyloxy methyl ketones and synthesis of (±)-actinopolymorphol B. J. Org. Chem. 2011, 76 (2), 500–511.

(4) Spanier, L.; Ciglia, E.; Hansen, F. K.; Kuna, K.; Frank, W.; Gohlke, H.; Kurz, T. Design, synthesis, and conformational analysis of trispyrimidonamides as α-helix mimetics. J. Org. Chem. 2014, 79 (4), 1582–1593.

